# OCT4 interprets and enhances nucleosome flexibility

**DOI:** 10.1101/2021.04.27.441583

**Authors:** Caitlin M. MacCarthy, Jan Huertas, Claudia Ortmeier, Hermann vom Bruch, Deike Reinke, Astrid Sander, Tim Bergbrede, Hans R. Schöler, Vlad Cojocaru

## Abstract

Pioneer transcription factors are proteins that induce cellular identity transitions by binding to inaccessible regions of DNA in nuclear chromatin. They contribute to chromatin opening and recruit other factors to regulatory DNA elements. The structural features and dynamics modulating their interaction with nucleosomes are still unresolved. From a combination of experiments and molecular simulations, we reveal here how the pioneer factor and master regulator of pluripotency, Oct4, interprets and enhances nucleosome structural flexibility. The magnitude of Oct4’s impact on nucleosome dynamics depends on the binding site position and the mobility of the unstructured tails of nucleosomal histone proteins. Oct4 propagates and stabilizes open nucleosome conformations by specific sequence recognition and nonspecific DNA exploration. Our findings provide a structural basis for the versatility of transcription factors in engaging with nucleosomes and have implications for understanding how pioneer factors induce chromatin dynamics.

## 1 Introduction

In eukaryotic cells, the DNA containing all the information about the cell’s fate and function is packed into chromatin inside the nucleus. The nucleosome, the basic unit of chromatin, is a nucleoprotein complex formed by ≈ 145-147 base pairs of DNA wrapped around an octamer of four histones proteins (H3, H4, H2A and H2B). Histones have a globular domain and disordered, charged regions at the N-terminus (and at the C-terminus of histone H2A), known as the histone tails [1].

The reading and regulation of this genetic information is carried out by site-specific DNA binding proteins called transcription factors (TFs) [2]. The extent of genome packing leaves a sizeable fraction of the DNA inaccessible for the majority of TF binding due to occlusion of binding sites by the histone core. Interestingly, a subgroup of TFs known as pioneer TFs (pTFs) can recognize and bind their motifs in nucleosome occupied regions of the genome [3, 4, 5]. A recent systematic study revealed that several families of transcription factors can have pioneer activity, binding with different affinities and mechanisms to nucleosomes [6].

One particularly interesting pTF is Oct4, a master regulator of pluripotency [7]. Oct4 cooperates with pTFs Sox2 and Klf4 in the conversion of somatic cells to pluripotent stem cells during reprogramming [8]. Structurally, Oct4 contains a DNA binding domain divided into two subdomains, the POU specific domain (POU_S_) and the POU homeodomain (POU_HD_), connected by a flexible linker. Together, the two subdomains recognize an octamer sequence in free DNA [9]. Research on Oct4 pTF activity reported that in nucleosomes Oct4 recognizes only half of its eight base pair binding site due to the twist of DNA and steric clashes with the histone core, which inhibit the canonical binding of the second subdomain [10, 11, 12, 13, 14].

Oct4 is known to bind cooperatively with Sox2 in the genome [15, 16, 17, 18, 19, 20, 21, 22]. Recent studies suggest they may also work together as pioneers on nucleosomal DNA [23, 10, 11], where one TF binding event alters DNA-histone contacts and DNA positioning, facilitating the binding of the second TF [22, 13, 14, 24]. However, characterizing the molecular mechanism that controls the binding of pTFs to the nucleosome, or the effect this binding has on nucleosome dynamics (and chromatin structure) remains an open challenge [5].

Recently, structures of pTFs bound to a nucleosome have been reported. The first published structure suggested a mechanism by which Sox2 binds and unravels nucleosomes [25]. The second revealed Oct4 together with Sox2 bound to a nucleosome [13]. Neither structure gave a complete picture: in the first structure, the entire outer gyres of the nucleosomal DNA were not resolved, whereas in the second structure only the Oct4 POU_S_ was resolved. The histone tails were not resolved in either of these structures. These and many other structural studies of pTFs use synthetic sequences, which are proven to tightly bind the histone core, making them well-suited for structural techniques, but not giving the full representation of nucleosome behavior *in vivo.*

In addition, recent studies resolved the structure of a genomic nucleosome that can be bound by Oct4 [12, 26]. While they did not present the structure of the Oct4-nucleosome complex, they served to reveal a novel Oct4 binding site not previously described by Soufi et al. [11]. Echigoya et al. speculated that the POU_S_ subdomain of Oct4 recognizes this binding site[12]. Moreover, they also showed that Oct4 interacts with the histone H3 and competes with the linker histone for binding. Both this and the work of Michael et al. [13] highlighted the importance of the POU_S_ but not the POU_HD_ subdomain in Oct4’s nucleosome binding. On the other hand, from molecular simulations using a reduced representation (coarse-grained) of the molecular species, Tan et al. also observed binding of Oct4 to this novel site [14]. However, the simulations revealed that the POU_HD_ domain recognizes the site. This is also supported by the DNA sequence that bares a typical homeodomain binding site.

We previously used molecular modeling and short molecular dynamics (MD) simulations to build models of Oct4 bound to nucleosomes on the binding sites proposed by Soufi et al. [10, 11]. From these, we confirmed that the canonical configuration of Oct4 with the subdomains bound on opposite faces of DNA is incompatible with nucleosome binding and we proposed alternative binding modes [27].

Here we report the structural features and dynamics involved in the Oct4-nucleosome interaction at atomic resolution. We used a combination of in *vitro* experiments and atomistic MD simulations to understand how Oct4 binds to its motifs in two genomic nucleosomes from regulatory sequences of the LIN28B and the ESRRB genes. Both sequences were chosen from the earliest data set demonstrating Oct4’s pioneer function [10] for their importance in defining stem cell pluripotency [28, 29, 30, 31] and their distinct Oct4 binding sites. We demonstrate Oct4’s sequence specific binding to the two native nucleosomes, and confirm that Oct4 favors binding sites positioned near the DNA ends. We show that Oct4 recognizes its binding sites using either DNA binding subdomain, and requires some nucleosome structural flexibility for efficient binding. Finally, we describe how Oct4 binding impacts the structural flexibility of the nucleosomal DNA at atomistic resolution. We found that the magnitude of this impact depends on the position of the binding site, the mobility of histone tails, and the motion of the Oct4 DNA binding subdomains.

## 2 Materials and Methods

### 2.1 Full-length Oct4 expression and purification

Experiments were all performed with purified full-length Oct4 from *M. musculus.* Briefly, Oct4 was cloned into the pOPIN expression vector using the SLIC method and Phusion Flash High-Fidelity PCR Master Mix (Finnzymes/New England Biolabs). SLIC reactions were then transformed into One Shot™ OmniMAC™ 2 T1^®^ Chemically Competent *E. coli* (ThermoFisher Scientific; Waltham, MA). After sequencing, the pOPIN-cHis-Oct4 construct was co-transfected with flashBACULTRA™ bacmid DNA (Oxford Expression Technologies; Oxford, UK) into Sf9 cells (ThermoFisher Scientific) using Cellfectin II^®^ (ThermoFisher Scientific) to generate recombinant baculovirus. Mid-log phase Sf9 cells were used to amplify the virus. Suspension High Five™ cells were infected with P3 virus for two days at 27 *°* C and 120 rpm shaking. After expression, crude lysates were purified on a HiTtap TALON column (GE Healthcare; Chicago, IL), cleaved on the column with 3C protease and followed by size exclusion chromatography (HiLoad Superdex 200, GE Healthcare). The final product was collected in 25 mM HEPES pH 7.8, 150 mM NaCl, 1 mM TCEP, and 5% glycerol with around 95% purity confirmed by SDS-PAGE.

### 2.2 Nucleosome reconstitution

The LIN28B and ESRRB nucleosome sequences were taken from Soufi et al [10] (Hg18 chr6:105, 638,004-105,638,165 and Hg18 chr14:75,995,474-75,995,636, respectively, see also Supplementary Methods). Sequences were selected for their role in pluripotency [29, 30] and their RNA induction after 48 hours of reprogramming. WT and mutant sequences were purchased from IDT (Coralville, IA, USA) with flanking AvaI restriction sites, sequenced, amplified in *E. coli,* digested, and finally purified by native PAGE using electroelution. After Cy5 labeling, DNAs were reconstituted at DNA:octamer ratios ranging from 1:1.2-1:1.6 with purified full-length *D. melanogaster* histone octamer using the salt-gradient dialysis method previously described [32], final buffer composition: 10 mM HEPES pH 7.6, 50 mM NaCl, 1 mM EDTA, and 0.5 mM DTT. Following dialysis, nucleosomes were heat shifted at 37°C for 2 hours and then checked for quality and concentration by native PAGE. Histone stoichiometry was checked by 22% SDS-PAGE followed by coomassie (R-250) (SERVA, Heidelberg, Germany) staining.

### 2.3 Electrophoretic mobility shifts and competition assays

For binding reactions, 20 nM nucleosomes were incubated with 0.05-0.4 μM of purified Oct4 in binding buffer (25 mM HEPES pH 7.6, 50 mM NaCl, 0.5 mM EDTA, 1 μg/μL BSA, 0.8 mM DTT, and 10% glycerol) for 1 hour at 25 °C. After incubation, reactions were run directly on a 6% native polyacrylamide gel (acrylamide/bis-, 37.5/1) containing 27 mM Tris-borate, 0.6 mM EDTA, and 5% glycerol and run in the same buffer. All Cy5-labeled DNA was detected using Fujifilm FLA-9000 (GE Healthcare). Competition assays using specific and nonspecific oligos were performed as previously described [11] using 0.2-4 μM of competitor, 20 nM nucleosome, and 105 nM Oct4. Oct4 off-rates from nucleosomes were determined by incubating specific competitor with the pre-formed Oct4/nucleosome complex that occurs after 1 hour incubation at 25 °C. Off-rate conditions were empirically determined for LIN28B and ESRRB nucleosomes due to the substantial differences in complex stability: LIN28B – 9 nM nucleosome, 45 nM Oct4, and 0.2-3.5 μM unlabeled competitor at 25 °C for 30 minutes; and ESRRB – 9 nM nucleosome, 67.5 nM Oct4, and 9-90 nM unlabeled competitor at 25 °C for 5 minutes. A short oligo containing the ESRRB Oct4 binding sequence was used as competitor for the ESRRB nucleosomes: AAGTGATAGTTATGCAGAGCGAATGGAGGG. For LIN28B, the specific competitor sequence published in Soufi et al. was used [11]. Experiments were performed in triplicate and densitometry was carried out using a DNA standard curve and Quantity One^®^ software (Bio-Rad, Hercules, CA). The values reported in Figure 2 were calculated by dividing the portion of Oct4-nucleosome complex by the value of starting free nucleosome (control). Two-way ANOVA using Sidak’s multiple comparison test was performed on relevant mean and standard deviation values. Statistics were done using Prism 7.0a for Mac.

### 2.4 Crosslinking experiments

Assemblies were generated as described in a previous section. Half of the assembly preparation was incubated with a final concentration of 1% formaldehyde on ice for 15 minutes. Crosslinking reactions were quenched by adding glycine to 250 mM final concentration. Crosslinking efficiency was checked by incubating an aliquot of crosslinked and uncrosslinked nucleosome in 15 mM MgCl_2_ and 300 mM NaCl at 60 ° C for 15 minutes and then running the samples on a 6% native PA gel. Samples were purified on a 10–30% sucrose gradient spun at 30,000 x g and 4 ° C in a Beckman Coulter Optima L-100 XP swing bucket rotor (SW-41; Brea, CA) for 18 hours. Fractions were collected from the bottom of the gradient, screened on native gels, pooled, and quantitated by densitometry using a DNA standard curve and Quantity One^®^ software (Bio-Rad, Hercules, CA). Off-rates with crosslinked nucleosomes were performed as described in the previous section.

### 2.5 Modelling Oct4-nucleosome complexes

To build the structural models of the LIN28B and ESRRB nucleosomes, we first selected 168 base pair sequences from the data by Soufi et al. [10, 11]. The selection was based on a comparison of ChIP-Seq data (accession code GSE36570) for Oct4 binding during reprogramming of fibroblasts to pluripotency (48 h after induction of Oct4 and the other three transcription factors required) with MNase-seq data revealing nucleosome positioning in human fibroblasts (accession code GSM543311) [33]. We considered the region around the MNase peak to correspond to the dyad and optimized the accessibility of the Oct4 binding sites proposed by Soufi et al. (S^-1.5^ and HD^-4.5^ on LIN28B and S^+5.5^ on ESRRB) [10, 11] (see Supplementary Methods). Then we threaded these selected sequences on a 168 base pairs nucleosome with the original Widom 601 DNA sequence [34] and with *Drosophila* or human histones by swapping each base according to the new sequence. For this, we used the “swapna” function in Chimera [35]. The complete 168 base pair Widom nucleosomes were built using the DNA from the 3LZ0 structure with the histones (including the histone tails) modelled using the 2PYO and 1KX5 structures as templates (see Supplementary Methods). We previously performed extensive MD simulations with these nucleosomes [36].

Next, we modelled Oct4-nucleosome complexes using the *Drosophila* LIN28B nucleosome and human ESRRB nucleosome models (see Supplementary Methods). The initial configuration of Oct4 we took from the following structures: (i) Oct4 bound to DNA in the canonical configuration with the two subdomains bound on opposite sides of DNA [37] that were built based on the structure of Oct4 bound as a homodimer to the PORE motif [38]. (ii) Oct4 bound with both subdomains on the same side of DNA (MORE configuration) obtained from the structure of Oct4 bound as the homodimer to the MORE motif [39] by stripping one monomer. (iii) Oct4 configurations obtained from a 100 ns MD simulation of apo Oct4 [27] performed with the same protocol as described below.

To model the binding of Oct4 to the nucleosomes, the binding sites from the structures of Oct4 bound to free DNA were superposed to the binding sites on the nucleosomes, using the base pairs that form sequencespecific interactions with Oct4. Then the free DNA was removed. The Oct4 configurations from the MD simulation of apo Oct4 were superposed on the initial models of canonical Oct4-nucleosome complexes [27] by fitting the subdomain that binds specifically to the DNA. Then, we removed the canonical configuration of Oct4. The resulting models of Oct4-nucleosome complexes had no steric clashes between the nonspecifically bound subdomain and the core histones. All models were validated in initial 100 ns classical MD simulations (protocol below). The validating short simulations of Oct4 bound to the LIN28B nucleosome on the binding sites proposed by Soufi et al. (HD^-4.5^ and S^-1.5^) were presented in our previous study [27].

### 2.6 Molecular dynamics simulations

Classical MD (cMD) simulations were performed as previously described [27, 36]. Every simulated species was first solvated in a truncated octahedron box of SPC/E water molecules, with a layer of at least 12 Å of water around the solute. Na^+^ ions were added to counter the negative charges of the system. K^+^ and Cl^-^ ions were added, up to a concentration of 150 mM. The systems were then optimized with an energy minimization, performed with the AMBER software[40]. Then, the systems were equilibrated for 13.5 ns, using *NAMD* [41]. The equilibration protocol was adapted from Jerabek et al. [39] and is described in detail in the Supplementary Methods. Harmonic distance restraints were applied to maintain DNA base pairing and Oct4-DNA base interactions. The force constant for these was gradually decreased. At the latest stages, the equilibration was unrestrained. Then, we performed production simulations in *NAMD*, in the isobaric-isothermic (NPT, p = 1 atm, T = 300K) ensemble, with Langevin dynamics for temperature control and a Nosé-Hoover and Langevin piston for pressure control. The Li-Merz ion parameters [42], the ff14SB [43] and the parmbsc1 force fields [44] were used for ions, protein, and DNA, respectively. Each individual simulation was 1 or 2 *μ*s long and multiple replicas were performed (Table 1).

**Table 1:** Overview of the simulations performed.

| Simulation name<sup>#</sup> | DNA | Binding site | Oct4 configuration | Time | $R_g$<sup>##</sup> | Number of Oct4 - DNA contacts<sup>###</sup> | | | |
| --- | --- | --- | --- | --- | --- | --- | --- | --- | --- |
| | | | | | | $\text{POU}_{\text{S}}$ | | $\text{POU}_{\text{HD}}$ | |
|  |  |  |  |  |  | Bases | Backbone | Bases | Backbone |
| LIN28B<sub>1</sub> | LIN28B | - | - | 1 $\mu\text{s}$ | 48.34-50.90 | - | - | - | - |
| LIN28B<sub>2</sub> | LIN28B | - | - | 1 $\mu\text{s}$ | 48.00-49.91 | - | - | - | - |
| ESRRB<sub>1</sub> | ESRRB | - | - | 1 $\mu\text{s}$ | 47.83-51.32 | - | - | - | - |
| ESRRB<sub>1-b</sub> | ESRRB | - | - | 1 $\mu\text{s}$ | 48.82-52.45 | - | - | - | - |
| ESRRB<sub>2</sub> | ESRRB | - | - | 1 $\mu\text{s}$ | 47.14-48.18 | - | - | - | - |
| $\text{HD}^{-7}\text{rev}_1$ | LIN28B | $\text{HD}^{-7}$ | Canonical | 1 $\mu\text{s}$ | 49.30-50.35 | 5-28 | 27-99 | 86-126 | 172-250 |
| $\text{HD}^{-7}\text{rev}_2$ | LIN28B | $\text{HD}^{-7}$ | MD | 1 $\mu\text{s}$ | 49.58-50.60 | 0-10 | 0-73 | 36-81 | 90-172 |
| $\text{HD}^{-7}\text{rev}_3$ | LIN28B | $\text{HD}^{-7}$ | Canonical | 1 $\mu\text{s}$ | 49.64-50.89 | 33-109 | 63-155 | 83-131 | 147-254 |
| $\text{HD}^{-7}\text{rev}_4$ | LIN28B | $\text{HD}^{-7}$ | MD | 1 $\mu\text{s}$ | 48.57-49.84 | 2-14 | 23-104 | 40-87 | 132-212 |
| $\text{S}^{+5.5}_1$ | ESRRB | $\text{S}^{+5.5}$ | MD1 | 2 $\mu\text{s}$ | 52.04-56.64 | 101-135 | 132-236 | 10-41 | 62-179 |
| $\text{S}^{+5.5}_1$ - Oct4 | ESRRB | - | - | 2 $\mu\text{s}$ | 55.60-58.98 | - | - | - | - |
| $\text{S}^{+5.5}_2$ | ESRRB | $\text{S}^{+5.5}$ | MD2 | 2 $\mu\text{s}$ | 48.44-49.96 | 77-118 | 80-151 | 20-65 | 26-93 |
| $\text{S}^{+5.5}_{2-b}$ | ESRRB | $\text{S}^{+5.5}$ | After 1 $\mu\text{s}$ of $\text{S}^{+5.5}_2$ | 1 $\mu\text{s}$ | 51.29-57.59 | 80-108 | 136-187 | 36-91 | 22-78 |
| $\text{S}^{+5.5}_{2-b}$ - Oct4 | ESRRB | $\text{S}^{+5.5}$ | - | 1 $\mu\text{s}$ | 50.10-52.93 | - | - | - | - |
| $\text{S}^{+5.5}_3$ | ESRRB | $\text{S}^{+5.5}$ | MD1 | 2 $\mu\text{s}$ | 47.40-48.88 | 99-148 | 93-191 | 7-35 | 63-131 |
| $\text{S}^{+5.5}_4$ | ESRRB | $\text{S}^{+5.5}$ | MD2 | 1 $\mu\text{s}$ | 48.15-49.25 | 55-112 | 104-170 | 3-86 | 40-112 |
| $\text{S}^{+5.5}_5$ | ESRRB | $\text{S}^{+5.5}$ | MORE | 1 $\mu\text{s}$ | 47.60-48.87 | 82-111 | 81-146 | 12-64 | 57-190 |
| $\text{S}^{+5.5}_{\text{T1}}$ | ESRRB | $\text{S}^{+5.5}$ | MD1 | 2 $\mu\text{s}$ | 50.00-56.82 | 77-142 | 105-187 | 0-24 | 27-76 |
| $\text{S}^{+5.5}_{\text{T1}}$ - Oct4 | ESRRB | - | - | 1 $\mu\text{s}$ | 47.54-48.66 | - | - | - | - |
| $\text{S}^{+5.5}_{\text{T1-b1}}$ | ESRRB | $\text{S}^{+5.5}$ | After 800ns of $\text{S}^{+5.5}_{\text{T1}}$ | 250 ns | 54.89-58.23 | 73-124 | 112-183 | 0-15 | 0-70 |
| $\text{S}^{+5.5}_{\text{T2}}$ | ESRRB | $\text{S}^{+5.5}$ | After 250ns of $\text{S}^{+5.5}_{\text{T1-b1}}$ | 1 $\mu\text{s}$ | 52.63-55.13 | 74-104 | 104-161 | 28-84 | 108-177 |
| $\text{S}^{+5.5}_{\text{T1-b2}}$ | ESRRB | $\text{S}^{+5.5}$ | After 800ns of $\text{S}^{+5.5}_{\text{T1}}$ | 250 ns | 54.23-58.32 | 80-127 | 104-170 | 0-18 | 0-76 |
| $\text{S}^{+5.5}_{\text{T3}}$ | ESRRB | $\text{S}^{+5.5}$ | After 200ns of $\text{S}^{+5.5}_{\text{T1-b2}}$ | 1 $\mu\text{s}$ | 53.17-55.62 | 85-130 | 125-208 | 4-56 | 83-180 |

### 2.7 Biased molecular dynamics simulations

We used minimal inter-atomic distances and coordination numbers (*δ*_min_ and *C*) as collective variables to perform biased MD (bMD) simulations. These are defined in the *Colvar* module [45] implemented in *NAMD* and *VMD* [46] (https://colvars.github.io/). *δ*_mln_ between two groups of atoms were measured using the weighted mean distance collective variable *distanceInv* defined as follows:

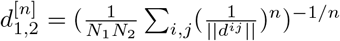

where ||*d^ij^*|| is the distance between atoms i and *j* in groups 1 and 2 respectively, and *n* is an even integer. This distance will asymptotically approach the minimal distance when increasing *n*. We used *n* =100. *C* between two groups of atoms were measured as:

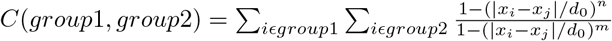

where *x_i_*, *x_j_* are the atomic coordinates of atoms *i* and *j*, *d*_0_ is the threshold distance (4.0 A), *n* and *m* are exponents that control the properties of the function for which we used the default values n=6, *m*=12.

The atom groups were defined using the Cα and P atoms from the proteins and DNA, respectively. To frequently sample nucleosome conformations in which the histone tails do not interact with the linker DNA, we applied harmonic wall biases to keep *δ*_min_ larger than 12 Å and *C* equal to 0 between the H3 and H2AC tails and the outer DNA gyre.

To sample the motion of the POU_HD_ between the two DNA gyres, we applied additional steered MD harmonic biases in which *δ*_min_ between POU_HD_ and the inner and outer gyres were changed with constant velocity from 5 to 15 Å and from 32 to 12 Å respectively and the *δ*_min_ between POU_HD_ and the 3’ L-DNA (last 11 base pairs) between 58 to 28 Å over 250 ns of simulation. To prevent the rapid closing of the nucleosome before the POU_HD_ moved between the gyres, near the L-DNA, we also applied the harmonic wall biases to keep *δ*_min_ and *C* between the H3 and H2AC tails and the outer DNA gyre larger than 30 Å and equal to 0, respectively, and to keep the *δ*_min_ between the two gyres larger than 20 A.

For all harmonic biases we used a force constant of 10 kcal/mol*Å^2^. After each biased simulation we performed additional, 1 *μ*s long classical simulations with or without Oct4 to ensure that the application of the biases did not distort dynamics.

### 2.8 Analysis of the MD simulations

All simulations were fitted to the histone core (excluding histone tails). Histone tails were defined as residues 1-45 for H3 (Human and *Drosophila*), 1-32 for H4 (Human and *Drosophila*), 1-18 and 119-129 for Human H2A, 1-17 and 116-124 for *Drosophila* H2A, 1-33 for Human H2B and 1-31 for *Drosophila* H2B. For the DNA, the 146 basepairs centered on the dyad were defined as the nucleosome core, whereas the remaining 11 basepair on each side were named as linker DNAs (L-DNAs)

The characterization of the breathing motions was done using the coordinate system originally described by Ozturk et al. [47], also used in our previous work [27, 36]. First, a coordinate system XYZ was defined, with the origin on the dyad. X was described as the vector along the dyad axis, Y as the cross product between X and a vector perpendicular to X intersecting it approximately at the center of the nucleosome, and Z as the cross product between X and Y. Two vectors, v_3_ and v_5_ were defined along the 3’ and 5’ L-DNAs. Then, the angle *γ*_1_ was defined as the angle between the projection of these vectors in the XZ plane and the Z axis, and *γ*_2_, as the angle between the projection of the vectors on the XY plane and the Y axis.

The mass-weighted radius of gyration of the DNA (R_g_) was calculated using the *cpptraj* software [48]. The same software was used for calculating the histone tail-DNA contacts and Oct4-DNA contacts. A contact was defined as a protein and a DNA non-hydrogen atoms closer than 4.5 A. Contacts were considered stable if they were present for more than 75% of the simulation.

## 3 Results

### 3.1 Oct4 binds to specific sites on nucleosomes with either DNA binding subdomain

First, we were curious about how each subdomain of Oct4 contributes to Oct4 binding at different locations of the two selected nucleosomes. To address this, we characterized purified Oct4 (Figure S1A) binding to a series of native and mutated nucleosomal DNAs (Figure 1). The native sequences we used depict the diversity of Oct4 genomic binding in that one sequence, LIN28B [11], contains multiple sites for Oct4 (Figure 1A) and other pTFs, whereas the other, ESRRB (Figure 1B) has a single Oct4 binding site. On LIN28B, the binding sites are located either towards the 5’ side (using the genomic 5’-3’ orientation as reference) or in the core of the nucleosome (Figure 1C), whereas on ESRRB the binding site lies in the core but close to the 3’ side (Figure 1D).

**Figure 1:**
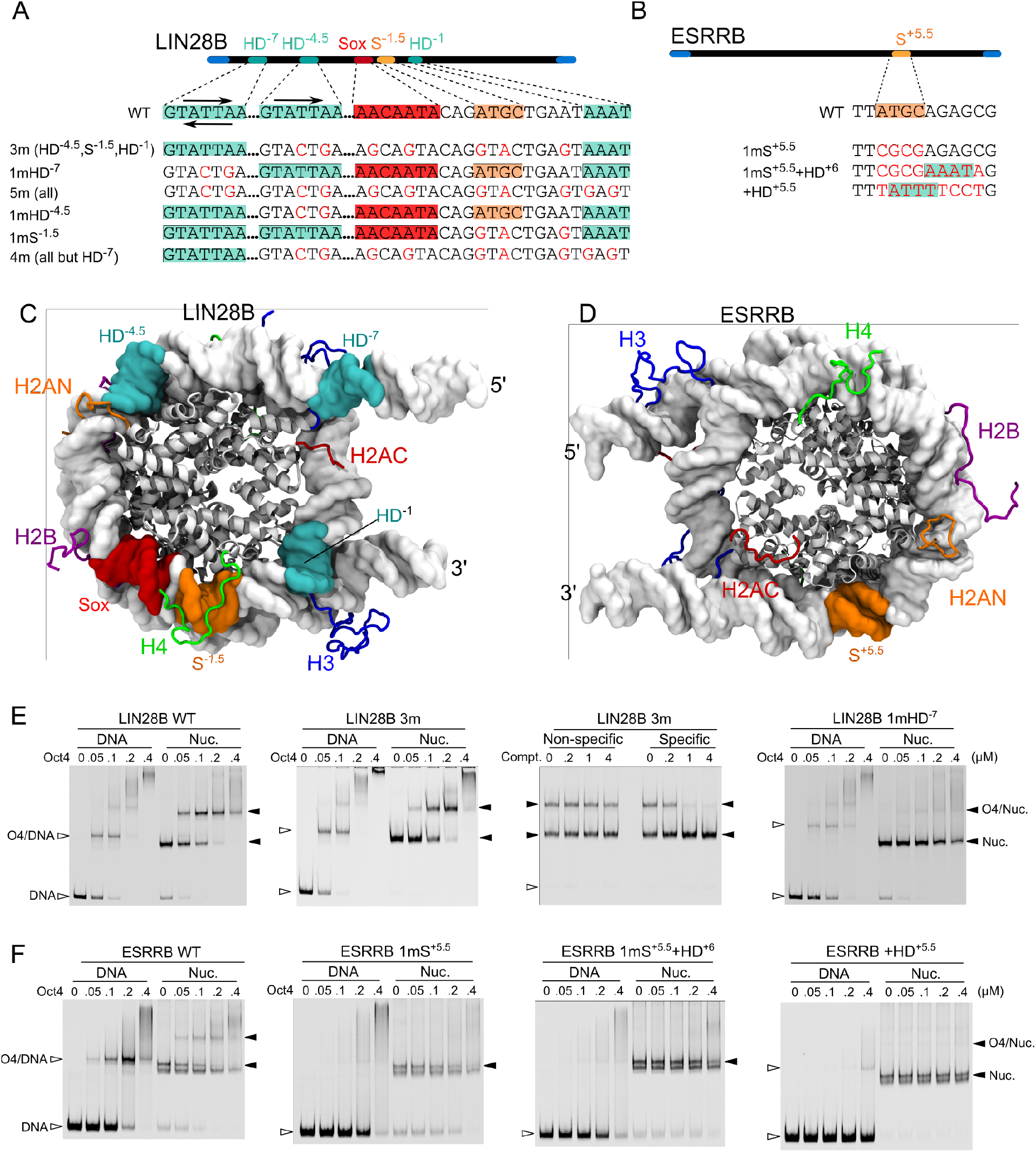
Oct4 binds to native nucleosomes at different positions with either DNA binding subdomain. (**A-B**) Schematics of the organization of the LIN28B (A) and ESRRB (B) genomic nucleosomes and mutants showing the positioning and sequences of the Oct4 binding sites. The mutants are named with a “m” preceded by the number of the binding sites mutated and followed by the names of these sites. For variants with multiple binding site mutations, the locations are shown in brackets. The POU_S_, POU_HD_, and Sox2 binding sites are in orange, teal, and red, respectively. Arrows indicate the binding orientation. The POUS binds in a 5’-ATGC-3’ orientation to free DNA, whereas the POU_HD_ in a 5’-T(A)AAT-3’ orientation with the N-terminal tail recognizing the first half and the globular part the second half. (**C-D**) Structural views of LIN28B (C) and ESRRB (D) with the histone core in gray cartoons, DNA in gray surface, binding sites in the corresponding colors, and histone tails in blue (H3), green (H4), orange (H2AN), red (H2AC), and purple (H2B). These representations and coloring are kept throughout the manuscript. (**E-F**) EMSAs of purified Oct4 with free DNA (left) and reconstituted nucleosomes (right). The third gel in 1E shows a competition assay using excess specific and nonspecific competitor. Filled horizontal arrowheads indicate nucleosomes or nucleosome-protein complexes and unfilled arrowheads, free DNA or DNA-protein complexes.

We first confirmed Oct4 binding to predicted nucleosomal sites by mutating key DNA bases within these sites and evaluating nucleosome binding in electrophoretic mobility shift assays (EMSAs). Assemblies were reconstituted using salt gradient dialysis with Cy5-labeled DNA and purified recombinant histones then checked for comparable positioning profiles (Figure S1B-I). Mutation of the Oct4 sites and the Sox site proposed by Soufi et al. (“3m”) [11] did not prevent Oct4 binding to LIN28B (Figure 1E). Given that Oct4 is also known to interact with histones [12, 49] or nonspecifically with DNA, we employed competition assays to test whether the persistent binding was specific or nonspecific. A molar excess of specific or nonspecific unlabeled double-stranded DNA was added to the Oct4-mutant nucleosome complex. Results showed that the interaction between Oct4 and the 3m nucleosome is sequence specific (Figure 1E, 3rd panel), suggesting the possibility of additional Oct4 binding sites on this nucleosome. A closer look at the LIN28B sequence revealed two potential homeodomain binding sites at superhelical locations (SHL) −7 and −1 (Figure 1A,C, HD^-7^ and HD^-1^). When we mutated all four Oct4 sites and the Sox site (5m), Oct4 nucleosome binding was reduced dramatically (Figure S2A,B).

In order to tease out the relative contributions of each site to overall Oct4-LIN28B binding, we mutated each site individually and in combinations leaving one site untouched. We observed that each site contributes to Oct4’s affinity, except for the Sox and HD^-1^ sites (Figures 1E, S2C,D). Mutating just the HD^-7^ site resulted in loss of Oct4 binding, suggesting that HD^-7^ is the primary Oct4 binding site (Figure 1E, 4th panel). While we performed the experiments and prepared the pre-print of this manuscript, two independent studies reported Oct4 binding to the HD^-7^ site, confirming our findings [12, 14]. Our results also indicate that the sites previously proposed by Soufi et al [10, 11] HD^-4.5^ and S^-1.5^ serve as secondary binding sites.

The HD^-7^ site contains a generic homeodomain binding site preceded by an adjacent POU_S_-like half-site (ATGA, not the canonical ATGC). Nevertheless, mutation of only the HD half-site reduced Oct4 binding drastically, demonstrating Oct4 POU_HD_-driven nucleosome binding (Figure 1E, 1mHD^-7^). Moreover, the sequence of the HD^-7^ and HD^-4.5^ are identical (Figure 1A), indicating Oct4’s preference for the HD^-7^ position and not the DNA sequence alone. Our data further suggests that Oct4 binds to LIN28B mainly through sequence specific binding of the POUHD to the HD^-7^ site.

Reconstitution of ESRRB WT and mutant sequences resulted in an assembly doublet (Figures 1F, S1E,F). One possible source of the doublet could be the eviction of one H2A/H2B dimer leading to a histone hexamer rather than an octamer in the nucleosome core [50]. Another possibility is that this sequence exists in two predominant conformations. Assemblies were routinely heat-shifted following salt-gradient dialysis suggesting both populations are thermodynamically stable (see Methods). Equal nucleosome-histone stoichiometry was also checked on coomassie stained SDS-polyacrylamide gels and histones appeared in comparable proportions, suggesting the presence of conformational isoforms (Figure S1E). Footprinting demonstrated not only were assemblies comparable between sequences but also suggested the presence of a middle positioned nucleosome (Figure S1G,H). Adding increasing amounts of free H2A/H2B dimer to histone octamer during assembly had no effect on the presence of either band, suggesting neither band contains a hexasome (Figure S1I).

In contrast to LIN28B, ChIP-Seq data show ESRRB contains one clear Oct4 binding peak at a canonical POU_S_ half-site followed by a potential POU_HD_ binding site (Figure 1B,D). To confirm this binding site, we first mutated the POU_S_ site, S^+5.5^, and observed a complete loss of Oct4 binding to both the nucleosome and free DNA (Figure 1F, 1st and 2nd panels, WT and 1mS^+5.5^). At high concentrations of Oct4, both free DNA and nucleosomes shifted to a smear rather than a defined band, which may be due to nonspecific interactions. Curious if a canonical homeodomain site would rescue binding, we integrated one directly 3’ to the POUS site, which did not restore binding to either free DNA or the nucleosome (Figure 1F, 3rd panel, 1mS^+5.5^+HD^+6^). This may be due to either the position of the POUHD facing into the histone core or that the homeoeodomain must work in concert with the POUS domain for stable binding. Interestingly, when we generated atomistic models of the ESRRB nucleosome we observed the POUS half-site facing away from the histone core suggesting it would be accessible for Oct4 binding (see later sections). To check the potential binding of the POU_HD_ on an exposed position at SHL +5.5, we introduced a canonical homeodomain half-site in the POU_S_ half-site location (Figure 1F, 4th panel, +HD^+5.5^). This mutation did not restore Oct4 binding to the nucleosome and displayed weak but distinct binding to free DNA. Together these results suggest that Oct4 binds to ESRRB at S^+5.5^ and the binding is driven by the POUS domain.

### 3.2 Oct4 uses nucleosome structural flexibility to bind

Recently, we reported differences in local nucleosome dynamics that extended beyond the linker DNA arms into the body of the nucleosome [27, 36]. Notably, dynamics were sequence dependent and greater in native nucleosomes, compared to the synthetic Widom 601 positioning sequence [34]. We wanted to know how restricting these nucleosome dynamics would influence Oct4 binding.

To test this, we crosslinked nucleosomes using formaldehyde (Figure S3A) before performing binding assays. Densitometry was performed on EMSA gels and presented as the ratio of Oct4-nucleosome complex relative to the amount of control nucleosome for each condition (Figure 2A-D). For LIN28B, crosslinking moderately increased binding at low Oct4 concentrations, but reduced binding at high concentrations (Figure 2A,B). Crosslinking ESRRB resulted in overall reduced Oct4 binding (Figure 2C,D). This suggests that the intrinsic dynamics of ESRRB facilitate Oct4 binding, while binding to LIN28B is more complex due to the presence of multiple Oct4 binding sites.

**Figure 2:**
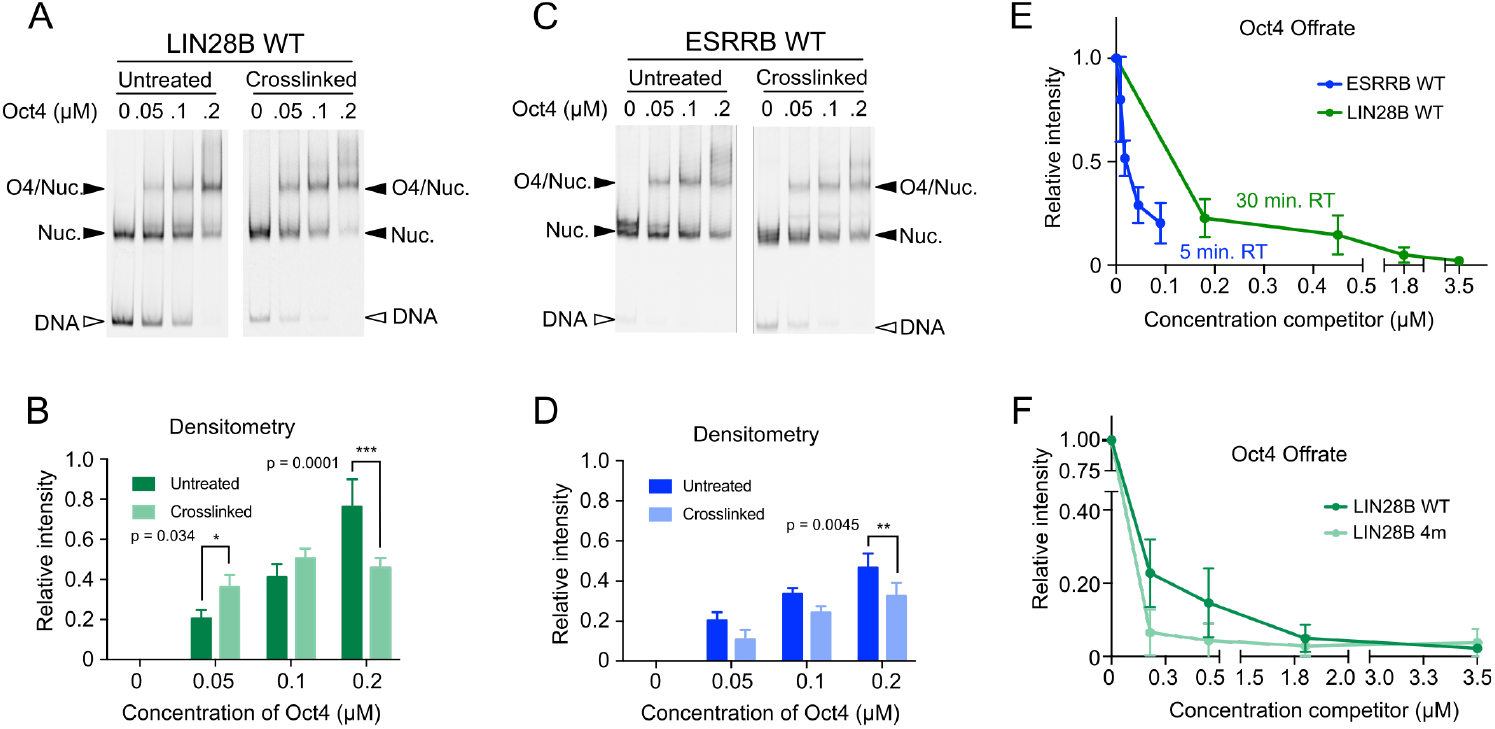
Nucleosome dynamics influence Oct4 binding. (**A and C**) EMSAs of purified Oct4 with untreated (left) or crosslinked (right) LIN28B (A) or ESRRB (C) nucleosomes. Filled arrowheads signal nucleosomes or nucleosome-protein complexes and unfilled arrowheads point to free DNA. (**B and D**) Bar graphs display the mean and standard deviation of densitometry relative to starting nucleosome amounts. n=3 per condition, * p=0.0344,** p=0.0045, *** p=0.0001. (**E**) Densitometry of Oct4 off-rate EMSAs using LIN28B (green) or ESRRB (blue) WT nucleosomes. Values represent the mean and the error bars reflect the standard of deviation. ESRRB n=4 and LIN28B n=5. (**F**) Same experiment as in panel E but comparing LIN28B WT and 4m nucleosomes (for 4m n=6).

We then wanted to know how the different sequences or mutants affected the stability of the Oct4-nucleosome complexes. After protein-nucleosome complex formation, we added increasing amounts of unlabeled specific competitor and monitored Oct4’s dissociation from the nucleosome. Notably, Oct4 separation from the LIN28B nucleosome required 20-40 fold higher amounts of competitor and six times longer incubation than from the ESRRB nucleosome (Figures 2E, S3B). These results corresponded with the number of Oct4 binding sites for each sequence, the higher affinity of the POU_HD_ to free DNA comparing to POU_S_ [37], and the established dynamics of the free nucleosomes [27]

When we mutated all Oct4 and Oct4-like binding sites except the primary HD^-7^ site (“4m”), the complex dissociated faster than the complex with the wild type LIN28B (Figure 2F, Figure S2C). This shows that although the mutated sites are not primary Oct4 binding sites, they do significantly influence the affinity of Oct4 for LIN28B. In summary, we observed that Oct4 has a higher affinity for the LIN28B nucleosome in part due to the multiple binding sites and the dynamics between the DNA and the histone core. Oct4’s affinity for the lone ESRRB nucleosome binding site is partially dependent on nucleosome dynamics and substantially lower than that for LIN28B.

### 3.3 Oct4 modifies nucleosome structural flexibility

One important recurring question about the function of pTFs is how their binding to nucleosomes leads to chromatin opening. To evaluate how Oct4 binding impacts on nucleosome dynamics, we generated structural models of Oct4 bound to the different binding sites (Figures 3, S4), performed MD simulations with the different models (Table 1), and compared them with the published simulations of free nucleosomes [27, 36]. We quantified nucleosome dynamics by measuring two features: overall nucleosome compactness using the radius of gyration (R_g_) and the breathing motions in two orthogonal planes, the plane of the core histones in a 2D top view of the nucleosome (angle *γ*_1_) and a plane perpendicular to it (angle *γ*_2_) [27, 36, 47] (see Methods).

**Figure 3:**
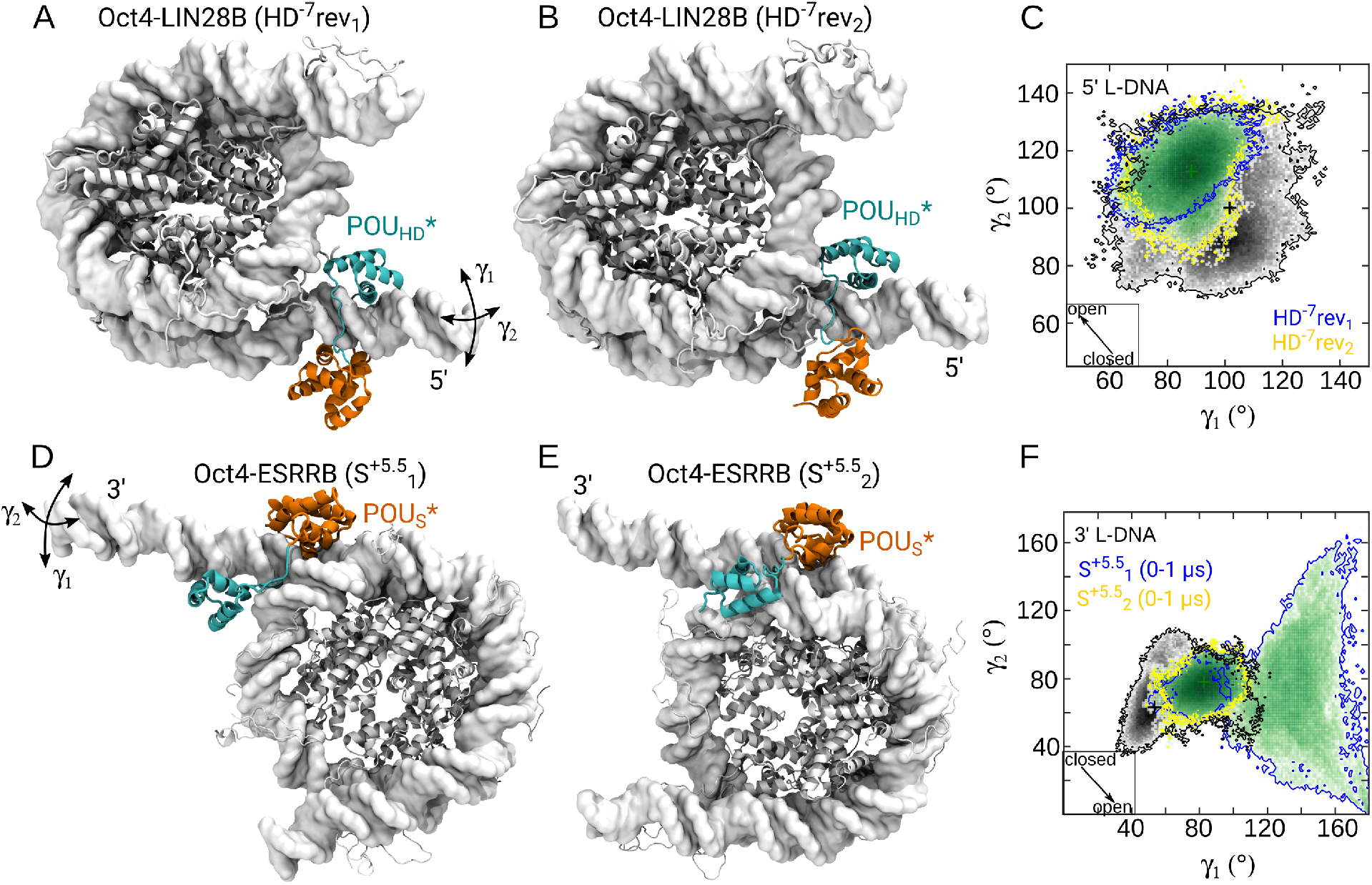
Oct4 binding modifies nucleosome breathing. (**A-B**) Representative structures from the simulations of the Oct4-LIN28B complex with Oct4 bound to the HD^-7^ binding site, in reverse orientation. The configuration of Oct4 is either canonical (A) or generated from a simulation of apo Oct4 (B) (see Methods). (**C**) 2D histogram depicting the γ_1_/γ_2_ conformational sampling in two independent simulations of the structures shown in A and B. (**D-E**) Representative structures from the simulations of the Oct4-ESRRB complex with Oct4 bound to the S^+5.5^ binding. Both starting configurations of Oct4 were taken from the simulation of apo Oct4. (**F**) 2D histogram depicting the γ_1_/γ_2_ conformational sampling in two independent simulations of the structures shown in D and E. The motions described by γ_1_ and γ_2_ are indicated with double headed arrows. In the histograms, in black is the sampling from 2 *μ*s of simulations of free nucleosome, whereas green marks the sampling from 2μs of simulations of Oct4-Nucleosome complexes. The sampling from each individual 1μs simulation of the complexes is in yellow and blue, respectively. The arrows in the square insets indicate the direction of the nucleosome opening. The “*” labels the sequence specific bound subdomain (also in subsequent figures)

We extended the simulations of Oct4 bound to the sites proposed by Soufi et al. [11] on LIN28B [27] to 1μs. Oct4 remained stably bound on this time-scale (Table 1) and did not modify the nucleosome dynamics (Figure S4A,B).

To model Oct4 bound to the HD^-7^ site, we used conformations of LIN28B partially open at the 5’ L-DNA from the simulations of the free nucleosome. In this way, we avoided clashes between the POU_HD_ domain of Oct4 and the other DNA gyre. We also modelled two binding orientations because the sequence is identical to that in the HD^-4-5^ site and we previously reported a preference for an atypical binding orientation [27]. The first has the POU_HD_ bound to the typical homeodomain site TAAT(AC), referred to as the ‘‘reverse” orientation because it’s on the 3’-5’ genomic DNA strand. The second orientation has the POU_HD_ bound to the GTAT(TA) motif on the genomic 5’-3’ DNA strand and is referred to as the ‘forward” orientation (Tables 1, S1). On both motifs the globular region of the POU_HD_ recognizes specifically the central AT bases. In our simulations of Oct4 bound to the HD^*4.5^ site which has an identical sequence, Oct4 remained stably bound only in forward orientation [27]. On the HD^-7^ binding site, Oct4 was stably bound in both orientations (Videos S1, S2). From here on, we only present the data for the reverse orientation (Figures 3A-C). The data for the forward orientation is in the Supplementary Material (Table S1, Figure S4B-E).

Due to the location of the HD^-7^ site in the L-DNA (Figure 1), Oct4 binding in the canonical configuration with the two subdomains embracing the DNA was also possible (Table 1, Figure 3A). To sample alternative configurations with the POU_g_ exploring nonspecifically the DNA region around this binding site, we generated models of Oct4 bound in configurations selected from MD simulations of free Oct4 (Figure 3B). The binding of Oct4 was stable on the 1μs timescale, regardless of the modelled configuration.

From the simulations, we found that when bound to the HD^-7^ binding site, Oct4 stabilized the LIN28B nucleosome in partly open conformations, which were rarely sampled in the simulations of the free nucleosomes. Moreover, the closed conformations of LIN28B were not sampled with Oct4 bound (Figures 3C, S4) because the specifically bound POU_HD_ blocked the closing.

On ESRRB, the binding of Oct4 in the canonical configuration was not possible because the binding site is located in the core nucleosomal DNA. Instead, we modelled the Oct4-nucleosome interaction using alternative Oct4 configurations (Table 1, Figure 3D,E, see also Methods). The DNA readout of the POU_S_ domain was maintained in all simulations analysed.

In one simulation of the Oct4-ESRRB complex (S^+5.5^_1_) the nucleosome opened with an amplitude significantly larger than the breathing amplitude of free nucleosomes on the same time scale (Table 1, Figures 3F, S5, S6, Video S3). The 3’ L-DNA, next to the S^+5.5^ binding site, displayed large amplitude motions in both directions defined by the 7 angles (Figures 3F, S5A). Nevertheless, in the other simulations, the nucleosome conformations were similar to those observed in the simulations of the free nucleosome (Video S4).

### 3.4 Oct4’s impact on nucleosome dynamics depends on histone tails

Next, we wanted to know whether the differences between the simulations of the Oct4-ESRRB complex are due to the histone tails, which are known to regulate inter- and intranucleosome dynamics [51, 36]. Therefore, we characterized how Oct4 binding modifies the interaction of histone tails with the DNA. For this, we calculated the number of stable tail-DNA contacts (defined as contacts present in at least 75% of the simulation). We focused on those tails in proximity to the HD^-7^ and S^+5.5^ binding sites on LIN28B and ESRRB, respectively (H3, H2AC, H4, and H2B from one monomer) and compared the simulations with and without Oct4 bound.

We previously reported that the free LIN28B nucleosome opened when the H3 and H2AC tails established fewer interactions with the L-DNA [36]. In particular, the H3 tail established fewer stable contacts with the 5’ L-DNA and more stable interactions with the core DNA (Figure 4A-C) facilitating nucleosome opening.

**Figure 4:**
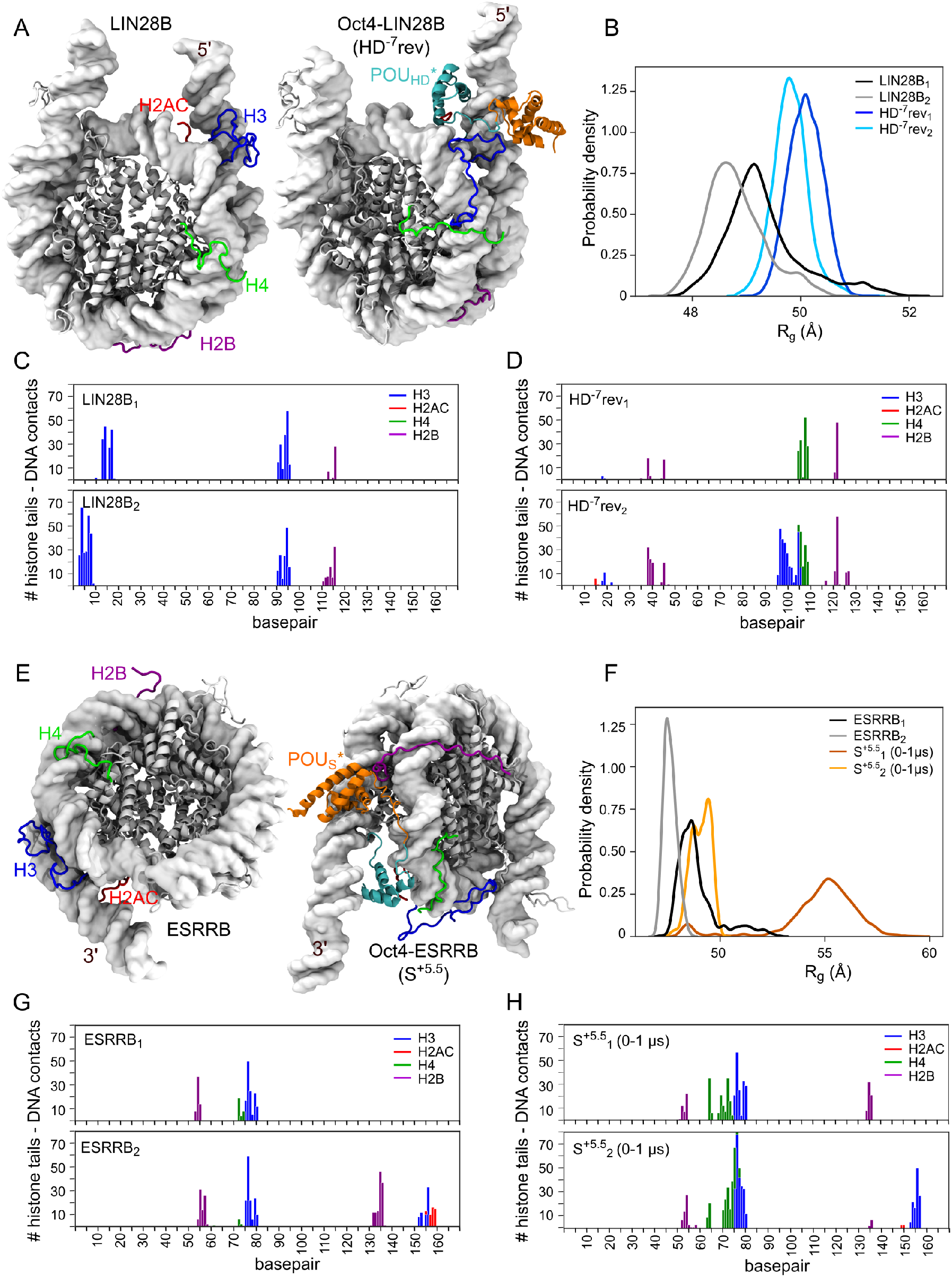
Oct4 modifies the histone tails-DNA interactions. (**A, E**) Representative structural snapshots of free nucleosome or Oct4-nucleosome complexes are shown together with the R_g_ histograms (B, F) to illustrate the nucleosome conformations sampled in the simulations for LIN28B and ESRRB, respectively. The Oct4-LIN28B complex (Oct4 bound to HD^-7^ in reverse orientation) and the Oct4-ESRRB complex (Oct4 bound to S+^5.5^). The histone tail – DNA interaction profiles show the number of stable contacts between the tails near the Oct4 binding site and the DNA for LIN28B (C,D) and ESRRB (G,H). A contact was defined as stable if it was present in more than the 75% of the 1μs simulation time.

When Oct4 was bound to the HD^-7^ site in the reverse orientation (Figure 4A,B,D, Figure S7), the nucleosome sampled mostly partially open conformations (Figure 4B) and almost all stable contacts of H3 to the 5’ L-DNA were lost (Figure 4C,D, Figure S7A,B). Access of the H3 tail to the L-DNA was blocked by the binding of the POUHD at the edge between the L-DNA and the core nucleosomal DNA (Figure 4A). The Oct4 linker in the canonical configuration (HD^-7^rev1 simulation) also blocked H3 tail binding near the dyad region (base pairs 90-95) (Figure 4D). The binding of Oct4 to the other sites proposed by Soufi et al [10] also resulted in fewer stable contacts between histone tails and the DNA (Figure S7C,D)

Both in the ESRRB alone and the Oct4-ESRRB complex (Figure 4E), the nucleosome remained closed (Figure 4F) in all simulations in which stable interactions were formed between the H3 and H2AC tails and the DNA at the 3’ end of the nucleosome (Figures 4G,H S8).

Independent of the DNA sequence and the location of the binding site, the number of stable H4 tail-DNA contacts increased upon Oct4 binding (Figures 4C,D,G,H, S7, S8).

When ESRRB remained closed with Oct4 bound, the H3 and H2AC tails interacted with both the inner and outer gyres of the DNA (Figure 5A,B). When ESRRB opened, these interactions were not formed (Figure 5C,D). Based on these findings, we hypothesized that the large amplitude opening occurred only in a single simulation of the Oct4-ESRRB complex because the required conformational transitions of the histone tails represent rare events on the 1 *μ*s timescale.

**Figure 5:**
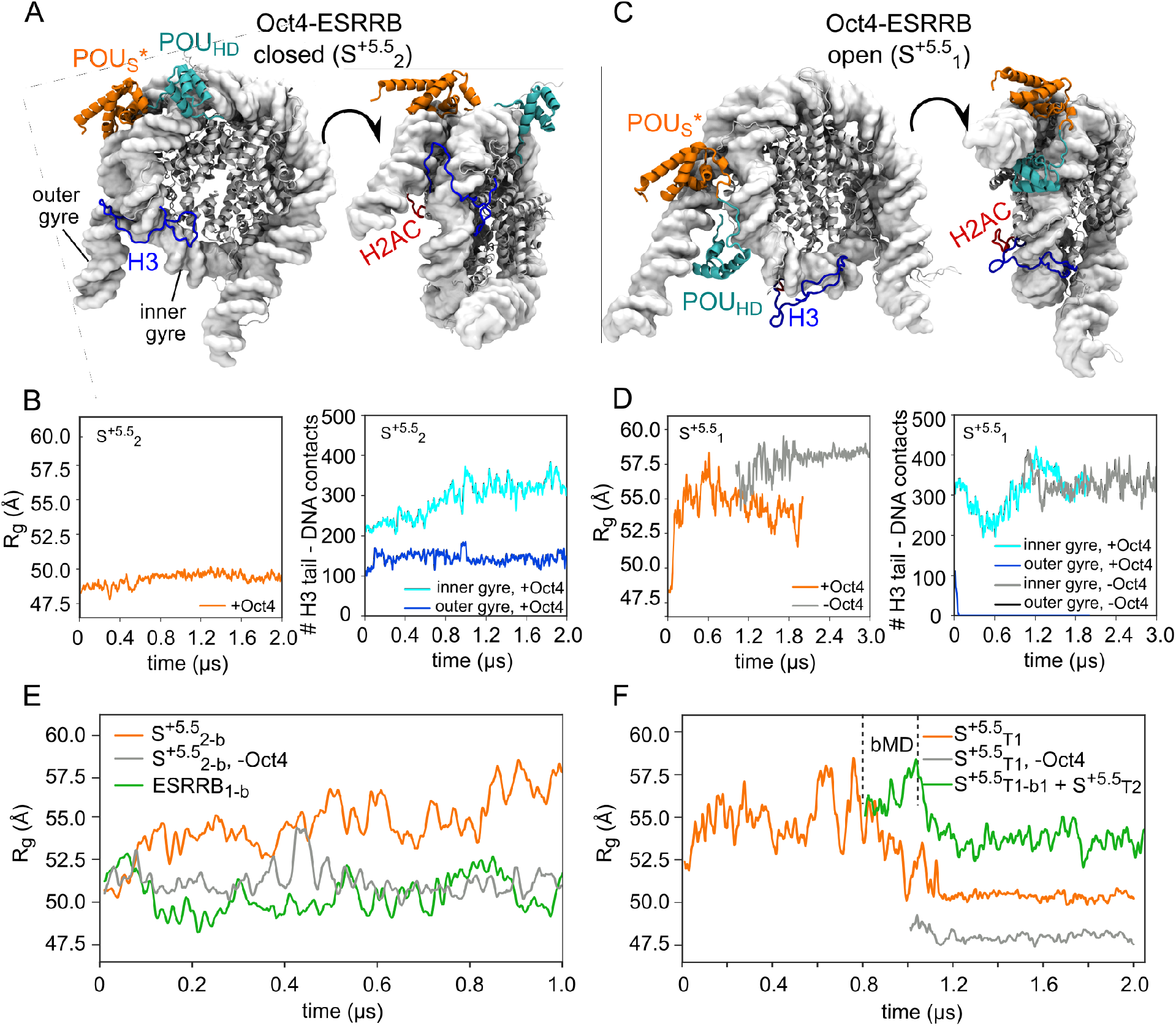
Oct4 stabilizes and enhances histone tail induced nucleosome opening. (**A**) Two views of a representative structure of the Oct4-ESRRB complex from the S^5.5^_2_ simulation in which the nucleosome remained closed. (**B**) Time series of R_g_ and the number of contacts between the H3 tail and the inner (middle 128 base pairs) and outer (terminal 20 base pairs) gyres of the DNA (in blue and cyan, respectively) from the S^5.5^_2_ simulation. (**C**) Same as (A) but from the S^5.5^_1_ simulation in which the nucleosome opened. (**D**) Same as (B) but from the S^5.5^_1_ simulation (orange) and from a simulation started after S^5.5^_1_ after removing Oct4 (gray). (**E**) Time series of R_g_ in biased MD simulations started from the S^5.5^_2_ simulation with (orange) or without Oct4 (gray) and from the ESRRB_1_ simulation (green). (**F**) Time series of R_g_ from three simulations started with the H3 and H2AC tails remodelled to adopt a configuration compatible with nucleosome opening: (S^5.5^_T1_, orange), a simulation (S^5.5^_T1_ -Oct4) started at 1 *μ*s of S^5.5^_T1_ after removing Oct4 (gray), and a simulation started after 800 ns of S^5.5^_T1_ in which the first 250 ns were biased (S^5.5^_T1-b1_) and the next 1 *μ*s were unbiased (S^5.5^_T2_, green). In S^5.5^_T1-b1_, the bias was applied to move the POU_HD_ in between the two gyres. The gray and green curves are drawn from the time snapshots of the S^5.5^_T1_ simulation (orange) at which the starting structures of the two simulations were selected (equilibrations are not shown).

To test this, we first performed biased MD simulations, in which we prevented the H3 and H2AC tails from forming interactions with the outer gyre of the DNA. For this, we added harmonic biases to the minimal tail-DNA interatomic distances and to the corresponding coordination numbers, which reflect the number of contacts (see Methods). We started two such simulations from two independent classical simulations of the Oct4-ESRRB complex in which the nucleosome remained closed (S^+5.5^_2_ and S^+5.5^_3_). In addition, we started two biased simulations from the same classical simulations after removing Oct4 and two biased simulations from the classical simulations of the ESRRB alone (Tables 1, S1, Figures 5E, S9A-C). ESRRB opened in all simulations but the amplitude of the opening was larger when Oct4 was bound (Figures 5E, S9A, Video S5). The opening was stable in 1μs of classical simulation started after the biased simulations (Figure S9B,C). Moreover, the sampling of the γ_1_/γ_2_ conformational space and the histone tails – interaction profiles were similar to those in the initial simulation in which ESRRB opened (S^5.5^_1_) (Figures S5B-D, S8B,C). These data indicates that the bias did not alter the natural dynamics of the systems.

Then, we also performed a classical simulation of the Oct4-ESRRB complex starting from a partially closed nucleosome but with the H3 and H2AC tails in configurations compatible with nucleosome opening (simulation S+^5.5^_T1_ (Table 1). Again, we observed a large amplitude opening of the nucleosome, similar to the one occurring in the original simulation (Figures 5F, S6A, S8D Video S6). However, after 1μs, the nucleosome closed into a compact conformation (Figure 5F, orange). This conformation was even more compact in a simulation started after 1μs of S^+5.5^_T1_ after removing Oct4 (Figure 5F, gray). Taken together, these data suggest a mechanism in which Oct4 stabilizes and further enhances the opening of ESRRB induced by specific H3 and H2AC tail configurations.

### 3.5 Oct4 stabilizes open nucleosomes by specific recognition and nonspecific exploration of DNA

Comparing the two classical simulations of the Oct4-ESRRB complex in which ESRRB opened extensively (S^+5.5^_1_ and S^+5.5^_T1_), we observed that in the first the open conformation was not only stable for 2*μ*s in the presence of Oct4 but it also remained stable when Oct4 was removed (Figures 5D, S5A). In the second, the nucleosome closed after 1μs (Figures 5F, S6A). Notably, in the first simulation the POUHD relocated between both DNA gyres by nonspecific exploration of the DNA. This prevented the nucleosome from closing (Figure 5C). To confirm whether the position of the POU_HD_ bridging the two gyres stabilized the open ESRRB conformation, we started two 250 ns biased simulations after 800 ns of the S^+5.5^T1 simulation (at maximum opening). In these, we forced the POUHD to move in between the gyres while the nucleosome remained open. For this, we used steered MD harmonic biases to change the minimial inter-atomic distances between the POUHD and the two gyres (see Methods). After 250 ns we switched off the bias and performed one additional classical simulation from each biased simulation (S^+5.5^_T2_, S^+5.5^_T3_, see Table 1). We found that in both these simulations, the nucleosome remained open for 1 *μ*s (Figures 5F, S9D, Video S7). Again, the sampling of the γ_1_/γ_2_ conformational space and the histone tails – interaction profiles were similar to those in the S^5.5^_1_ simulation (Figures S6B,C, S8D)

Visualizing the position of the Oct4 subdomains during 5 *μ*s aggregate simulation time of the Oct4-ESRRB complex (Figure 6A), we observed a narrow distribution for the position of the POU_S_ forming very contacts with the inner gyre due to its sequence specific binding to the outer gyre (Figure 6B). The position of the POU_HD_ showed a wider distribution with a large number of contacts both with DNA bases and the backbone in the inner gyre. When the ESRRB opened (simulation S^+5.5^_1_), its contacts with the inner gyre were maintained while new contacts were formed with the backbone of the outer gyre (Figure 6A). Similar distributions were observed in the 4 *μ*s simulations of the Oct4-ESRRB complex started with modified H3 and H2AC configurations (Figure 6C). In the first simulation (S^+5.5^_T1_) in which the nucleosome closed, the POU_HD_ explored only the inner gyre, whereas in the other two, it bridged the two gyres (Figure 6D).

**Figure 6:**
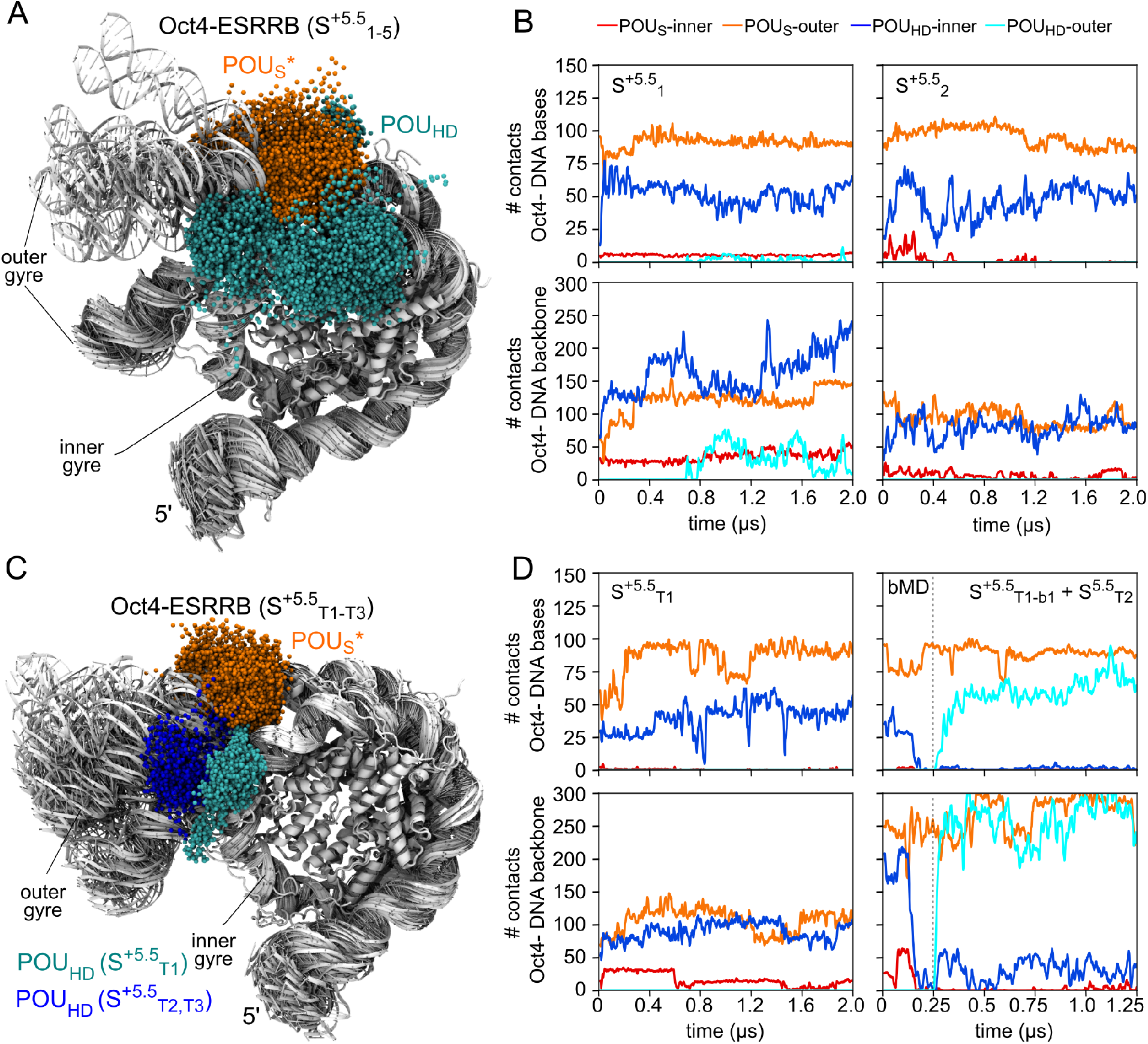
Oct4 modifies nucleosome dynamics by sequence specific recognition and nonspecific DNA exploration. (**A**) Nucleosome sampling by the Oct4 subdomains in 5 *μ*s aggregate simulations time of the Oct4-ESRRB complex. (**B**) Number of contacts between the POU_S_ (red, orange) or the POU_HD_ (blue, cyan) with DNA bases (sequence specific, upper plots) or DNA backbone (nonspecific, lower plots) of the inner gyre (red, blue) or the outer gyre (orange, cyan) in two of the simulations shown in (A) (S^+5.5^_1_ and S^+5.5^_2_). (**C**) Same as (A) in 4 *μ*s aggregate simulations time of the Oct4-ESRRB complex started with the H3 and H2AC tails remodelled to a configuration compatible with nucleosome opening. (**D**) Same as (B) but in the simulations shown in (C) (S^+5.5^_T1,T2,T3_).

On the other hand, the partially open conformation of the LIN28B nucleosome (Figure S10A) was stabilized by the sequence specific binding of the POU_HD_ to the outer gyre and by nonspecific interaction of the POU_S_ with the outer gyre. Very few contacts were formed between either subdomain and the inner gyre when Oct4 was bound on the HD^-7^ site of LIN28B (Figure S10B). There was no difference between Oct4 bound in reverse or forward orientations (Figure S10C,D).

## 4 Discussion

How pioneer factors [4, 5] contribute to the transition of closed, silent chromatin to transcriptionally active DNA remains poorly understood. They bind to DNA wrapped in nucleosomes sometimes using only partial binding motifs and a range of translational and orientation binding preferences, suggesting a diverse range of potential pTF mechanisms [6]. One mechanism proposes that for Oct4-Sox2 composite motifs, Sox2 binding to the minor groove deforms the DNA and mechanically loosens the grip of histones, freeing up buried binding sites and facilitating Oct4 binding [13, 25]. This mechanism involves Sox2-induced DNA opening and the use of only one of Oct4’s DNA binding subdomains, the POU_S_ domain. These structural studies provided the first evidence for a direct transcription factor-induced nucleosome opening. However, several questions remained unanswered: (i) Does the multi-domain pTF Oct4 alone have a similar impact on the nucleosome as Sox2?, (ii) How do pTFs bind native nucleosome sequences? and (iii) How do histone tails modulate or adapt to the binding?

Here we found that that Oct4 can use either domain to recognize its binding sites on nucleosomes and we describe at atomic resolution the interplay between intra-nucleosome dynamics and pTF binding (summarized in Figure 7).

**Figure 7:**
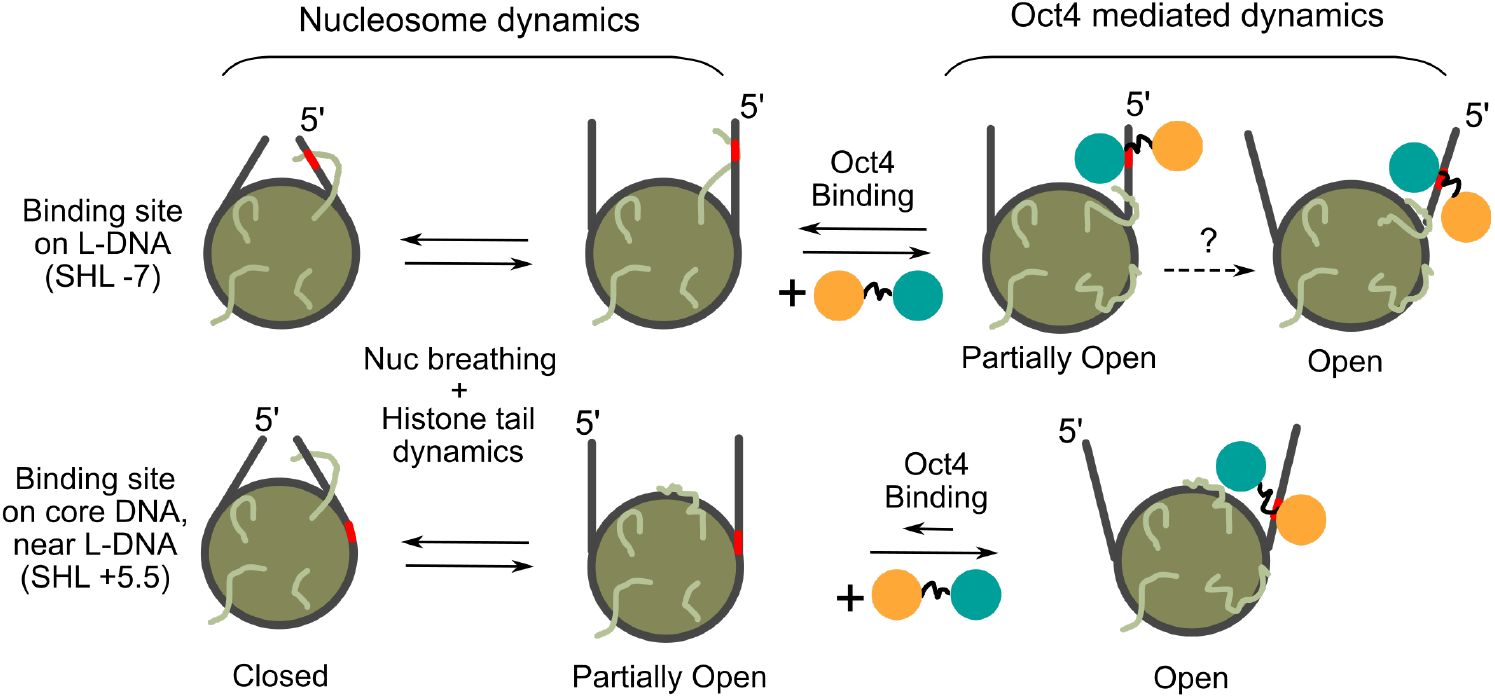
Summary of Oct4 mediated nucleosome dynamics. The Oct4 POUS and POUHD subdomains are shown as orange and teal circles, respectively. Larger circles with straight lines represent nucleosomes and their linker DNA as viewed from the top in closed, partially open, or fully open configurations. The red points on linker DNA indicate Oct4’s sequence specific binding half-sites. Coiled lines represent histone tails. Dotted arrows indicate hypothetical transitions.

Oct4 binds to one native nucleosome (LIN28B) primarily on a POU_HD_ binding site at the 5’ end of the nucleosomal DNA (the HD^-7^ site), but also to the POU_S_ and POU_HD_ sites located in the core nucleosomal DNA [11]. The binding to HD^-7^ was also proposed by two other studies while we were preparing the pre-print of this manuscript [12, 14]. We observed a lower affinity for the LIN28B WT nucleosome compared to the values reported by Soufi et al. [11]. This is most likely due to differences in experimental setup as well as the source of histones and Oct4 proteins (see Methods).

Mutating only the HD^-7^ site reduced Oct4 binding substantially, whereas mutating the other binding sites had only a moderate effect on binding kinetics. These findings are consistent with recently published studies of the LIN28B nucleosome [12, 11, 14]. Overall, these data point to Oct4’s adaptability as a pTF as it is able to recognize multiple binding sites on the LIN28B nucleosome using either the POU_S_ or POU_HD_.

On another native nucleosome (ESRRB), Oct4 binds to the POU_S_ binding site positioned at SHL +5.5. The dissociation of Oct4 from this nucleosome was significantly faster than dissociation from LIN28B. This may be explained both by the difference in the number of binding sites between the two nucleosomes and by the higher affinity of the POUHD domain to the DNA [9, 37]. Additionally, we showed that the binding of Oct4 to crosslinked nucleosomes is reduced at high concentrations of Oct4. The moderate increase in the binding of Oct4 to crosslinked LIN28B at low Oct4 concentrations can be explained by the location of the predominant HD^-7^ on the linker DNA arm. Our models and simulations show that the binding of Oct4 to this higher affinity site requires more open conformations of LIN28B, which LIN28B often adopts [27, 36]. When crosslinked, the LIN28B nucleosome HD^-7^ site is fixed in a range of conformations. At lower concentations, Oct4 could predominantly bind to more open conformations with higher affinity, which then become saturated at higher Oct4 concentrations. Without crosslinking, the intrinsic nucleosome dynamics facilitate Oct4 binding predominantly to partially open conformations. Another possibility is an induced fit mechanism in which Oct4 facilitates the opening of closed LIN28B conformations upon initial nonspecific binding. The other sites with lower Oct4 binding affinity located on the nucleosome core are only occupied at higher Oct4 concentrations. Therefore, we propose that Oct4 uses the intrinsic structural flexibility of the nucleosome to access its binding sites. And this is likely to be more important in genomic nucleosomes because they are generally believed to be more flexible than those assembled on engineered sequences [52, 53, 54].

To study how Oct4 impacts nucleosome dynamics we built structural models of Oct4-nucleosome complexes. First, we modelled free nucleosomes by positioning the dyad at the center of the reported micrococcal nuclease (MNase) peak [10, 27, 36]. This is an approximation as MNase-seq does not reveal the nucleosome position at base pair resolution. The recent cryo-EM structure of the free LIN28B, resolved after we completed our first set of simulations, differs from our models by 1 base pair [26]. This difference is minimal and can be attributed to the intrinsic flexibility displayed by LIN28B [27, 36]. Moreover, the structure was resolved with 147 and not 168 base pairs of DNA, difference that may also lead to minimal variations in positioning. Therefore, the approximation we used to position the nucleosomes at the time when no experimental structure was available is now validated. In our models, all confirmed Oct4 binding sites are exposed and accessible for the binding of the corresponding Oct4 subdomain we conclude that the models we used are reliable for studying the Oct4-nucleosome interaction.

To validate our models, we performed short MD simulations after we fixed Oct4-DNA contacts in the equilibration phase. The contacts were selected from the crystal structure of Oct4 bound to free DNA [38]. This assumed the same contacts are formed with the nucleosomes. This is now confirmed for the POU_g_ domain in the new structure of Oct4-Sox2-nucleosome complex [13]. For the HD^-4.5^ binding site, we previously tested two different binding orientations and found that the binding is stable only in one of them (forward) [27]. Since the sequence in the HD^-4.5^ and HD^-7^ sites are identical, we used the same approach to model the Oct4 binding to the HD^-7^ site and found that the complex was stable in both orientations. Therefore, the question whether the binding orientation of the POUHD is important for nucleosome recognition remains unanswered.

Regardless of the orientation, Oct4 can bind to the HD^-7^ site only if LIN28B is in a partially open conformation. Oct4 stabilized such conformations by the sequence specific binding of the POUHD, occupying the space between the core and linker DNA (Figure 7). The POU_S_ domain remained near the LIN28B DNA, sampling a range of positions. This suggests that one mechanism by which Oct4 impacts chromatin dynamics is by restricting the breathing of the nucleosome towards partly open nucleosome conformations. Although we did not observe an Oct4 induced opening of LIN28B, we can’t exclude this as a possibility due to the limited sampling achieved in our simulations. However, when bound to the binding sites located in the core nucleosomal DNA, Oct4 did not affect the dynamics of LIN28B. Nevertheless, given that LIN28B is bound by several pioneer transcription factors, it is also possible that the effect of Oct4 on some of the binding sites depends on the binding of Sox2 of Klf4. The experiments by Tan and Takada, reveal an allosteric mechanism by which the binding of Sox2 affects the posterior binding of Oct4 [14]. Overall, this suggests that the impact of Oct4 binding on nucleosome dynamics depends on the position of the binding site.

In contrast, we observed a large Oct4-mediated opening of the ESRRB nucleosome in one of the initial classical simulations of the Oct4-ESRRB complex. The open conformation was stable even after Oct4 was removed. We confirmed it is an Oct4-mediated opening using biased MD simulations. These findings suggest that under certain conditions, the binding of Oct4 induces and stabilizes open conformations. However, to do this, Oct4 first interprets the intrinsic nucleosome flexibility to bind to its sequence specific sites. For ESRRB, this was apparent when we showed that the free ESRRB nucleosome displayed the highest flexibility at the 3’ end of the nucleosomal DNA [36], precisely in the region where the Oct4 binding site is located.

Other pioneer transcription factors, such as Rap1 [55] and NF-κB [56] have also been shown to induce or stabilize open conformations of chromatin fibers or mononucleosomes configurations. Rap1 also uses intrinsic nucleosome and chromatin dynamics to facilitate binding [55]. Moreover, it is established that enhanced breathing of nucleosomes has an impact on the structure and dynamics of chromatin fibers [57, 58]. Therefore, the mechanism we describe for Oct4 may be general to pTF function and represents one way pTF binding modifies structural features in chromatin.

The major opening of ESRRB depended on two factors: (i) the position and conformation of the histone H3 and H2AC tails, and (ii) the motion of the POU_HD_ domain of Oct4, which explored the nucleosomal DNA in a nonspecific manner. The position and conformation of the H3 and H2AC tails are critical for nucleosome breathing [36, 59, 60]. We proposed that substantial, short lived opening of free nucleosomes is only possible when these tails establish a minimum number of interactions with the linker DNAs [36]. Here we also observed that the long lived Oct4 mediated nucleosome opening depends on the mobility of histone tails. As long as the tails embraced the L-DNA arms and bridged them to the core nucleosomal DNA, the opening did not occur. The H3 and H2AC tails also influenced Oct4 binding to the LIN28B nucleosome. When Oct4 was bound to the HD^-7^ site, both tails were displaced into a configuration with fewer contacts with the L-DNA arms. Taken together, these findings suggest that any transition to a pTF induced metastable open nucleosome conformation relies on the mobility of histone tails regulating intrinsic nucleosome breathing. Finally, in all our simulations, the binding of Oct4 to the nucleosome was accompanied by a substantial increase of the H4-nucleosome contacts. Because the H4 tail is essential to establish nucleosome-nucleosome interactions [61, 62, 63], the binding of Oct4 might also alter the higher order chromatin structures. The interplay between TF binding and histone tails was also observed in the Sox2 bound nucleosome structure in which the H4 tail was displaced compared to the free nucleosome [25].

In addition to the role of histone tails in any TF mediated nucleosome opening, our simulations revealed distinct roles for the Oct4 domains in nucleosome binding and dynamics. One domain is recognizing specific DNA sequences on nucleosomes while the other explores both DNA gyres in a sequence nonspecific manner. While exploring, the domains may get trapped in positions that prevent an open nucleosome from closing. How often this happens in the genomic context remains unclear. We observed one such stable position of the POUHD between the core and the linker DNA of the ESRRB nucleosome, which stabilized an extensively open nucleosome conformation that remained stable even when Oct4 was removed. This suggests that pTFs with multiple DNA binding domains might use their domains not only for sequence specific recognition of the binding site but also to establish barriers to nucleosome closing or inter-nucleosome stacking by binding sequentially distant pieces of DNA. The relative motion of the domains in one TF is restricted by the linker between them. This may further explain why a major effect on nucleosome dynamics was observed only for certain binding site positions. For example, on LIN28B, the binding sites located in the nucleosome core are too far from the linker DNA arm to allow bridging of the two gyres by the Oct4 domains.

Finally, our study is a demonstration of the strength of molecular simulations in revealing the structural features and the dynamic mechanisms involved in pTF-nucleosome binding. We previously showed that short MD simulations are sufficient to exclude Oct4 binding modes that are incompatible with nucleosome structure [27]. Here we show that MD simulations not only revealed pTF-induced conformational transitions in nucleosomes but also enabled us to discover their mechanisms at atomic resolution. However, all atom MD simulations are mainly limited by the amount of sampling achieved [64]. Despite reporting here extensive aggregate simulation times, the sampling achieved is still not sufficient to observe multiple opening events in a single long simulation. In addition, the accuracy of the force fields used for simulating the dynamics of unstructured regions such as the histone tails is still under scrutiny [65, 66]. Nevertheless, this approach has been successfully employed to investigate nucleosome dynamics and can be used to study the structure and dynamics of nucleosome-protein complexes for which no experimental structure is available [67].

## 5 Acknowledgements

We would like to thank Peter Becker and Catherine Regnard for their support by providing us with plasmids, reagents, technical assistance, and, most importantly, for their advice and active discussions. We would also like to thank Ken Zaret and Abdenour Soufi for providing us with alignments and native nucleosome sequences. Also, thank you to Francesca Mattiroli for guidance with the manuscript as well as the experimental approach. J.H. is part of the International Max Planck Research School-Molecular Biomedicine, Munster, Germany. This work was supported by funds of the Max Planck Society and The Royal Netherlands Academy of Arts and Sciences. The MD simulations were supported by the Gauss Centre for Supercomputing e.V. (www.gauss-centre.eu), project ID 12622, STRUCNUCREC running on the GCS Supercomputer SuperMUC at the Leibniz Supercomputing Centre (www.lrz.de).

## Notes

### Competing Interest Statement

The authors have declared no competing interest.

### Summary of Updates

In this revised manuscript, we include new data from extending some of the molecular dynamics simulations we reported in the first version and from a substantial amount of additional simulations. We also repeated some of the experiments presented in the first version and provide new explanations and validations of the experimental results. The new results further strengthen our conclusions about the interplay between Oct4 binding to nucleosomes and nucleosome structural flexibility. In addition, we improved the clarity and readability of the text.

