## Supporting Methods and Data for "OCT4 interprets and enhances nucleosome flexibility"

### Supplementary Material

### Supplementary Methods

#### DNaseI and restriction enzyme nucleosome footprinting

DNaseI footprinting - Cy5 3' single end-labeled nucleosomal DNAs were generated by PCR using the validated constructs from the section "Nucleosome reconstitution" as templates and then purified from 10% native gels by electroelution. Assemblies were reconstituted as described in the main methods section. Approximately 600 fmol of assembly or free DNA were partially digested with DNaseI (Thermo Fischer Scientific, USA) at 25°C for 2 minutes in EMSA binding buffer (25 mM HEPES pH 7.6, 50 mM NaCl, 0.5 mM EDTA, 1 $\mu$g/$\mu$L BSA, 0.8 mM DTT, and 10% glycerol) supplemented with $MnCl_2$. Reactions were stopped with the addition of an equal volume of Stop Buffer (2x - 90% deionized formamide, 0.1% bromophenol blue, 20 mM EDTA) and then heated to 95°C for two minutes. The entire reaction was run on 15% denaturing gels (1x TBE and 8 M urea) at 50 mA for 4 hours. Gels were imaged using a Fujifilm FLA-9000 fluorescence scanner (GE Healthcare).

Restriction digest footprinting - ESRRB free DNA and assemblies were digested overnight with equal amounts of AluI, HhaI, HinfI, and NcoI restriction enzymes (NEB, Ipswich, MA, USA) at 37°C. Reactions were then heated at 95°C for 10 min. to inactivate restriction enzymes as well as denature and dissociate histone proteins. Reactions were run on 20% native PA gels at 10 mA/gel for 3.5 hours and then imaged on Fujifilm FLA-9000 (GE Healthcare). The gels were then stained with Gel Red (Sigma Aldrich) and imaged on a UV transilluminator (VWR, USA).

#### Building the structural model of the 168 bp Widom nucleosomes

First, we built a 145 bp Widom nucleosome with full-length *Drosophila* histones, including tails. For this, we used Modeller (https://salilab.org/modeller) with the PDB structures 2PYO (*Drosophila* histones and human $\alpha$-satelite DNA), 3LZ0 (*Xenopus Laevis* histones and Widom DNA) and 1KX5 (*Xenopus Laevis* histones and human $\alpha$-satelite DNA) as templates. The DNA was transferred as a rigid body from the 3LZ0 structure, the core histones were modelled based on the 2PYO structure and the histone tails by homology modelling based on the 1KX5 structure (the only structure in which the tails are resolved). 100 models were generated using a "slow" optimization protocol followed by a "slow" MD-based refinement protocol. The model with the lowest discrete optimized energy was extended to 195 base pairs of DNA with the original Widom 601 sequence (*26*). First, the 3 base pairs at each end of each strand in the 3LZ0 structure were removed because they differ from the original Widom sequence. Then, fragments of 28 base pairs B-DNA generated with Nucleic Acid Builder, from Ambertools 18 (*46*) were added to each end based on a sequence alignment between the DNA sequence from the 3LZ0 structure and the original Widom sequence. To optimize the computer resources needed for the simulations, we kept only 168 bp DNA in our final model (11 and 12 bp L-DNAs at the 5'and 3' ends, respectively. To obtain this final model, we removed 13 and 14 base pairs from the 5' and 3' respectively from our 195 bp model. We used this model with *Drosophila* histones as template to built the equivalent model of a Widom nucleosome with human histones by homology modeling. Again, we built 100 models and selected the one with the lowest normalized DOPE score as defined in Modeller. These two models were then used as scaffolds to build the ESRRB and LIN28B nucleosomes as described in the main text.

#### Building structural models of Oct4-nucleosome complexes

For the models of the Oct4-LIN28B and Oct4-ESRRB complexes, we used the nucleosome with *Drosophila* and human histone, respectively. Our choice for the origin of the histones has a simple reason related to the chronology in which the experiments and the simulations were performed. First, we modelled and simulated the Oct4-LIN28B complex with *Drosophila* histones to match our initial experiments. Then, we modelled the Oct4-ESRRB complex with human histones because Soufi et al *(10,11)* performed their studies in human cells. While we performed experiments in parallel with both human and *Drosophila* histones, we found that the nucleosome assembly was more successful with the *Drosophila* histones. However, at this point the main set of the simulations of the Oct4-ESRRB complex with human histones were already in advanced stages. Therefore we continued the experiments with *Drosophila* histones and the simulations with the human histones. We expect the binding modes of Oct4 to nucleosomes to be independent on the histone sequences.
The sequences of the LIN28B and ESRRB used for the nucleosome models with the main binding sites ($HD^{-7}$, $HD^{-4.5}$, $S^{-1.5}$ in LIN28B and $S^{+5.5}$ in ESRRB) highlighted as in Figure 1 and the dyad in red are:

LIN28B (human genome assembly NCBI36 - hg18):
chromosome 6: 105,637,999-105,638,166

```
  1 AGTTAAGT GGTATTAA CATATCCT CAGTGGTG AGTATTAA CATGGAAC TTACTCCA
 57 ACAATACA GATGCTGA ATAAATGT AGTCTAAG TGAAGGAA GAAGGAAA GGTGGGAG
113 CTGCCATC ACTCAGAA TTGTCCAG CAGGGATT GTGCAAGC TTGTGAAT AAAGACAC
```

ESRRB (human genome assembly NCBI36 - hg18):
chromosome 14: 75,995,470-75,995,637

```
  1 ATCAGCAG GGAGAAGG AGCGCCTC CCCATGTG GGACCTGG AGAAACAG AGGGTGGA
 57 GGGAGCAT AGAGAGTC TGTTCTAA GCTGCAAA GCAAAGGC CTGGCGAC CTAGGAGA
113 CCATGGAG TTCCAGAA AGTGATAG TTATGCAG AGCGAATG GAGGGAAT CAGCACGC
```

The experimental data from where these were selected is available at GEO, with the accession codes GSE36570 (ChIP-Seq data) (*11*) and GSM543311 (MNase-Seq data) (*44*).

All models of the Oct4-nucleosome complexes presented in this manuscript were validated by monitoring the sequence specific and nonspecific interactions with of the subdomain of Oct4 that is bound specifically to each analyzed binding site. All the results we present here are from simulations in which the sequence specific contacts Oct4 with the nucleosomal DNA were preserved (see also (*19*)).

#### Equilibration protocol

The equilibration protocol was adapted from our previous simulations of Oct4 bound to the free DNA and is shown in the table below (see (*29*)).

| Step | Time | Restraint type | Force constant (kcal/mol·$A^2$) | Restrained Atoms/Bonds* |
|---|---|---|---|---|
| 1+ | 150 ps | Positional | 25 | Protein and DNA heavy atoms |
| 2 | 150 ps | Positional | 10 | Protein and DNA heavy atoms |
| 3 | 250 ps | Positional | 5 | Protein and DNA heavy atoms |
| 4 | 250 ps | Positional | 1 | Protein and DNA heavy atoms |
| 5 | 250 ps | Positional<br>Distance | 1<br>25 | Protein and DNA backbone<br>DNA basepairs<br>OCT4-DNA base interactions |
| 6 | 250 ps | Positional<br>Distance | 1<br>10 | Protein and DNA backbone<br>DNA basepairs<br>OCT4-DNA base interactions |
| 7 | 250 ps | Positional<br>Distance | 1<br>5 | Protein and DNA backbone<br>DNA basepairs<br>OCT4-DNA base interactions |
| 8 | 250 ps | Distance | 1 | DNA basepairs<br>OCT4-DNA base interactions |
| 9 | 250 ps | Distance | 0.5 | DNA basepairs<br>OCT4-DNA base interactions |
| 10 | 250 ps | Distance | 0.1 | DNA basepairs<br>OCT4-DNA base interactions |
| 11 | 250 ps | Distance | 0.05 | DNA basepairs<br>OCT4-DNA base interactions |
| 12 | 250 ps | Distance | 0.01 | DNA basepairs<br>OCT4-DNA base interactions |
| 13 | 1.5 ns | - | - | |
| 14 | 2.25 ns<br>(1.5 fs)# | - | - | |
| 15 | 7 ns<br>(2 fs)# | - | - | |

*Atoms for positional restraints, bonds for distance restraints.
+ The first step includes the heating from 20 to 300K, and it's done in the NVT ensemble. All the other steps are done in the NPT ensemble at 300K.
### The number in brackets indicates the timestep used in that step. For steps 1-13, a timestep of 1 fs is used. For the nucleosome alone systems, the protocol is identical, omitting the OCT4-DNA restraints.

#### Supplementary Data

- **Supplementary Table S1**: Overview of the simulations presented in the supplementary data.
- **Figure S1**: Characterization of *in vitro* reconstituted native nucleosomes (supplementary to data in Figures 1,2).
- **Figure S2**: Validation of Oct4 binding sites on nucleosomes (supplementary to data in Figure 1).
- **Figure S3**: Crosslinking and competition experiments (supplementary to data in Figure 2).
- **Figure S4**: Breathing motions of LIN28B in simulations of the Oct4-LIN28B complex with Oct4 bound to different sites (supplementary to data in Figure 3).
- **Figure S5**: Breathing motions of ESRRB in classical and biased simulations of the ESRRB alone and the Oct4-ESRRB complex (supplementary to data in Figure 3).
- **Figure S6**: Breathing motions of ESRRB in classical and biased simulations of the Oct4-ESRRB complex started with modified H3 and H2AC tail configurations (supplementary to data in Figure 3).
- **Figure S7**: Histone tail - DNA interaction profiles in the Oct4-LIN28B complex (supplementary to data in Figure 4).
- **Figure S8**: Histone tail - DNA interaction profiles in the Oct4-ESRRB complex (supplementary to data in Figure 4).
- **Figure S9**: Nucleosome opening in biased simulations followed by classical simulations of the Oct4-ESRRB complex (supplementary to data in Figure 5).
- **Figure S10**: Sequence specific binding and nonspecific DNA exploration on LIN28B by the two Oct4 subdomains (supplementary to data in Figure 6).
- **Video S1**: 1 $\mu$s simulation of Oct4 bound to the LIN28B nucleosome, in the "reverse" configuration ($HD^{-7}rev_1$). The starting Oct4 configuration was selected from a simulation of apo Oct4. Oct4 stabilizes a partially open nucleosome conformation.
- **Video S2**: 1 $\mu$s simulation of Oct4 bound to the LIN28B nucleosome in the "forward" configuration ($HD^{-7}for_1$). The starting Oct4 configuration was selected from a simulation of apo Oct4. Oct4 stabilizes a partially open nucleosome conformation
- **Video S3**: 2 $\mu$s simulation of Oct4 bound to the ESRRB nucleosome ($S^{+5.5}{}_1$). The starting Oct4 configuration was selected from a simulation of apo Oct4. Oct4 stabilizes an extensively open nucleosome conformation when the $POU_{HD}$ subdomain moves in between the two DNA gyres.
- **Video S4**: 1 $\mu$s simulation of Oct4 bound to the ESRRB nucleosome ($S^{+5.5}{}_2$). The starting Oct4 configuration was selected from a simulation of apo Oct4 (different than the one used in Video S3). The nucleosome remains closed due to the position of the unstructured tails of histones H3 and H2AC
- **Video S5**: Biased simulations of Oct4 bound to the ESRRB nucleosome. 1 $\mu$s of a simulation where histone tails H3 and H2AC are biased to move away from the outer DNA gyre($S^{+5.5}{}_{2\text{-b}}$), followed by 1$\mu$s of unbiased simulation ($S^{+5.5}{}_{2'}$). The starting configuration was the snapshot after 1$\mu$s of the $S^{+5.5}{}_2$ simulation. The nucleosome remains open since the histone tails don't reform the contacts with the outer DNA gyre.

- **Video S6**: 2 $\mu$s simulation of Oct4 bound to the ESRRB nucleosome ($S^{+5.5}{}_{T1}$). The starting Oct4 configuration was selected from a simulation of apo Oct4. The starting configurations of the H3 and H2AC tails were taken from the simulation ($S^{+5.5}{}_{1}$) in which the nuclesome opened. Oct4 stabilizes an extensively open nucleosome conformation. After 1 $\mu$s, the nucleosome closes because the $POU_{HD}$ subdomain of Oct4 which explores the DNA nonspecifically does not move into a position between the two DNA gyres and the interactions between the histone tails and the outer DNA gyre are reformed.

- **Video S7**: Biased simulations of Oct4 bound to the ESRRB nucleosome. 250 ns of a simulation where Oct4's $POU_{HD}$ is biased to move into the space between both gyres ($S^{+5.5}{}_{T1\text{-}b1}$), followed by 1$\mu$s of unbiased simulation ($S^{+5.5}{}_{T2}$). The starting configuration was the snapshot after 800ns of the $S^{+5.5}{}_{T1}$ simulation. The nucleosome remains open due to the position of Oct4's $POU_{HD}$ domain.

**Table S1: Overview of the simulations presented in the Supplementary data**

| Simulation name# | DNA | Binding site | Oct4 configuration | Time | $R_g$## | Number of Oct4 - DNA contacts### | | | |
|---|---|---|---|---|---|---|---|---|---|
| | | | | | | $POU_S$ | | $POU_{HD}$ | |
| | | | | | | Bases | Backbone | Bases | Backbone |
| $ESRRB_{2\text{-}b}$ | ESRRB | - | - | 1 $\mu$s | 50.10-52.93 | - | - | - | - |
| $HD^{-7}for_1$ | LIN28B | $HD^{-7}$ | Canonical | 1 $\mu$s | 49.16-51.51 | 25-56 | 69-117 | 92-131 | 177-253 |
| $HD^{-7}for_2$ | LIN28B | $HD^{-7}$ | MD | 1 $\mu$s | 49.11-50.08 | 0-26 | 10-68 | 60-117 | 116-188 |
| $HD^{-7}for_3$ | LIN28B | $HD^{-7}$ | Canonical | 1 $\mu$s | 49.17-51.11 | 35-59 | 74-143 | 62-121 | 150-220 |
| $HD^{-7}for_4$ | LIN28B | $HD^{-7}$ | MD | 1 $\mu$s | 49.24-51.85 | 3-56 | 32-127 | 42-86 | 57-139 |
| $S^{-1.5}{}_1$ | LIN28B | $S^{-1.5}$ | MORE | 1 $\mu$s | 48.38-49.54 | 69-102 | 64-97 | 6-39 | 50-125 |
| $S^{-1.5}{}_2$ | LIN28B | $S^{-1.5}$ | MD | 1 $\mu$s | 47.81-49.72 | 52-99 | 63-102 | 0-5 | 0-55 |
| $HD^{-4.5}for_1$ | LIN28B | $HD^{-4.5}$ | MORE | 1 $\mu$s | 48.97-50.83 | 12-72 | 72-127 | 35-61 | 68-105 |
| $HD^{-4.5}for_2$ | LIN28B | $HD^{-4.5}$ | MD | 1 $\mu$s | 47.88-49.28 | 0-8 | 3-48 | 71-104 | 117-192 |
| $S^{+5.5}{}_{3\text{-}b}$ | ESRRB | $S^{+5.5}$ | After 1 $\mu$s of $S^{+5.5}{}_3$ | 1 $\mu$s | 49.99-56.52 | 103-150 | 129-235 | 14-41 | 77-151 |
| $S^{+5.5}{}_{3\text{-}b}$ -Oct4 | ESRRB | $S^{+5.5}$ | After 1 $\mu$s of $S^{+5.5}{}_3$ | 1 $\mu$s | 50.78-55.64 | - | - | - | - |
| $S^{+5.5}{}_{2'}$ | ESRRB | $S^{+5.5}$ | After 1 $\mu$s of $S^{+5.5}{}_{2\text{-}b}$ | 1 $\mu$s | 53.28-57.99 | 73-99 | 133-172 | 56-104 | 53-137 |
| $S^{+5.5}{}_{2'}$ -Oct4 | ESRRB | $S^{+5.5}$ | After 1 $\mu$s of $S^{+5.5}{}_{2\text{-}b}$ | 1 $\mu$s | 55.08-59.64 | - | - | - | - |
| $S^{+5.5}{}_{3'}$ | ESRRB | $S^{+5.5}$ | After 1 $\mu$s of $S^{+5.5}{}_{3\text{-}b}$ | 1 $\mu$s | 54.52-56.62 | 73-98 | 146-189 | 38-61 | 151-225 |
| $S^{+5.5}{}_{3'}$ -Oct4 | ESRRB | $S^{+5.5}$ | After 1 $\mu$s of $S^{+5.5}{}_{3\text{-}b}$ | 1 $\mu$s | 54.77-57.75 | - | - | - | - |

### "T" labels simulations started with modified H3 and H2AC tail configurations; "b" labels biased simulations

#### $R_g$ and the number of contacts are shown as the percentiles 5 to 95.

##### A contact was defined as a non-hydrogen atom closer than 4.5 Å to another non-hydrogen atom.

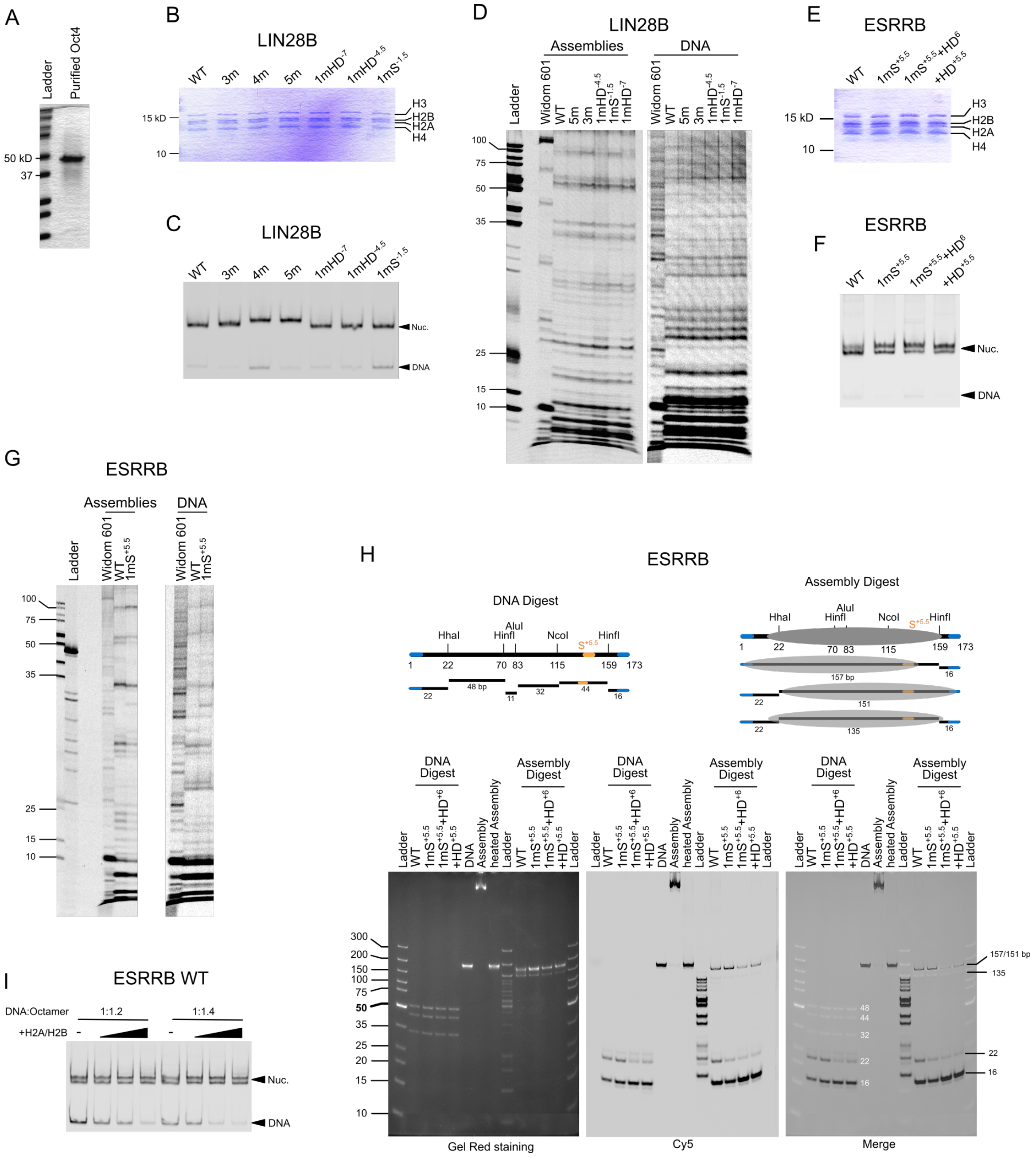


**Figure S1: Characterization of *in vitro* reconstituted native nucleosomes.** (**A, B, E**) Coomassie stained SDS-PA gels of (A) purified Oct4 protein (see main methods), (B) LIN28B nucleosomes, or (E) ESRRB nucleosomes. (**C, F**) Cy5 signal from native gels loaded with (C) LIN28B or (F) ESRRB nucleosomes. Arrowheads indicate assembled nucleosomes and free DNA. (**D, G**) DNase I footprints of (D) LIN28B or (G) ESRRB assemblies. (**H**) Gel red and Cy5 signals from ESRRB assembly restriction enzyme footprinting run on native gels. Top panels are schematic representations of free DNA or assembly digests. Blue and orange colors indicate Cy5 labeled ends and the $POU_S$ half binding site ($S^{+5.5}$), respectively. Gray ovals represent the histone core position. Fragment length is indicated below fragments and on the merged gel. **I)** Cy5 signal from native gel loaded with ESRRB WT assemblies at two DNA:Octamer ratios supplemented with increaseing amounts of purified H2A/H2B dimer. Arrowheads indicate assembled nucleosomes and free DNA.

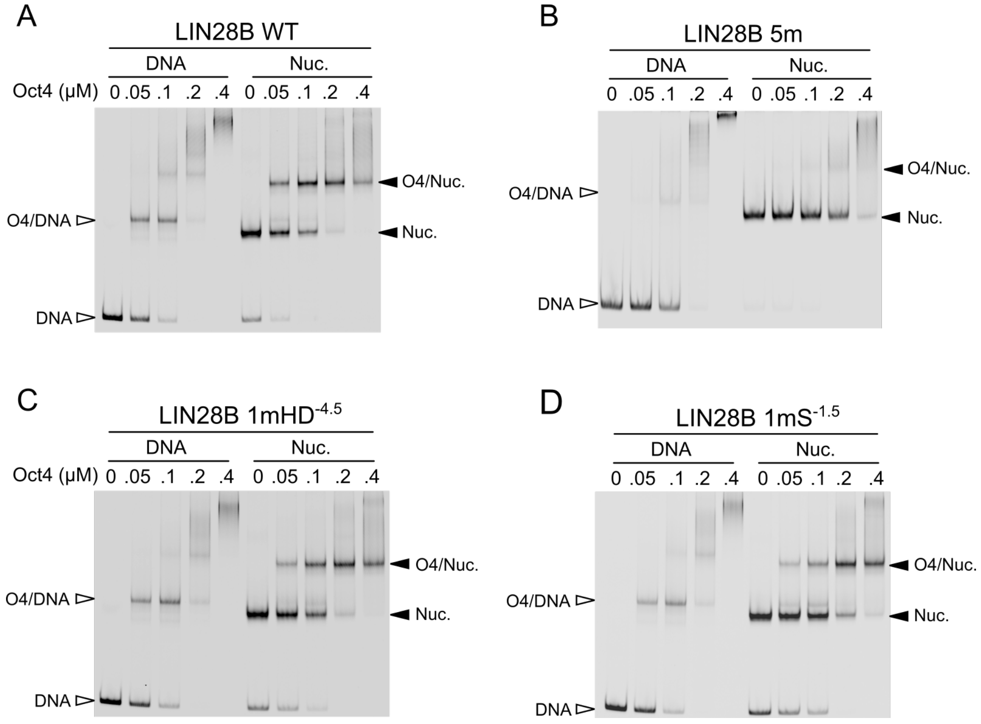


**Figure S2: Validation of Oct4 binding sites on nucleosomes.** EMSAs of purified Oct4 with free DNA (left) and reconstituted nucleosomes (right). Filled horizontal arrowheads indicate nucleosomes or nucleosome-protein complexes and unfilled ones, free DNA or DNA-protein complexes. Wild-type and mutant sequences can be found in main Figure 1.

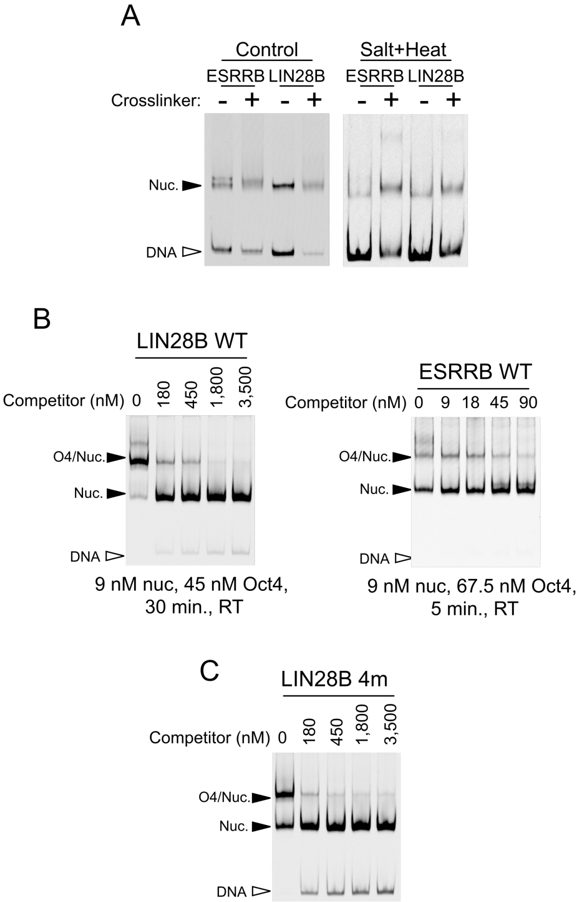


**Figure S3: Crosslinking and competition experiments.** (**A**) Native gels of LIN28B and ESRRB untreated and crosslinked nucleosomes. The right panel shows a representative native gel of a disassembly assay as a control for successful crosslinking. (**B-C**) Representative EMSAs of Oct4 off-rate experiments from Figure 2 panels E and F, respectively. Filled horizontal arrowheads indicate nucleosomes or nucleosome-protein complexes and unfilled ones, free DNA. See also Figure 2

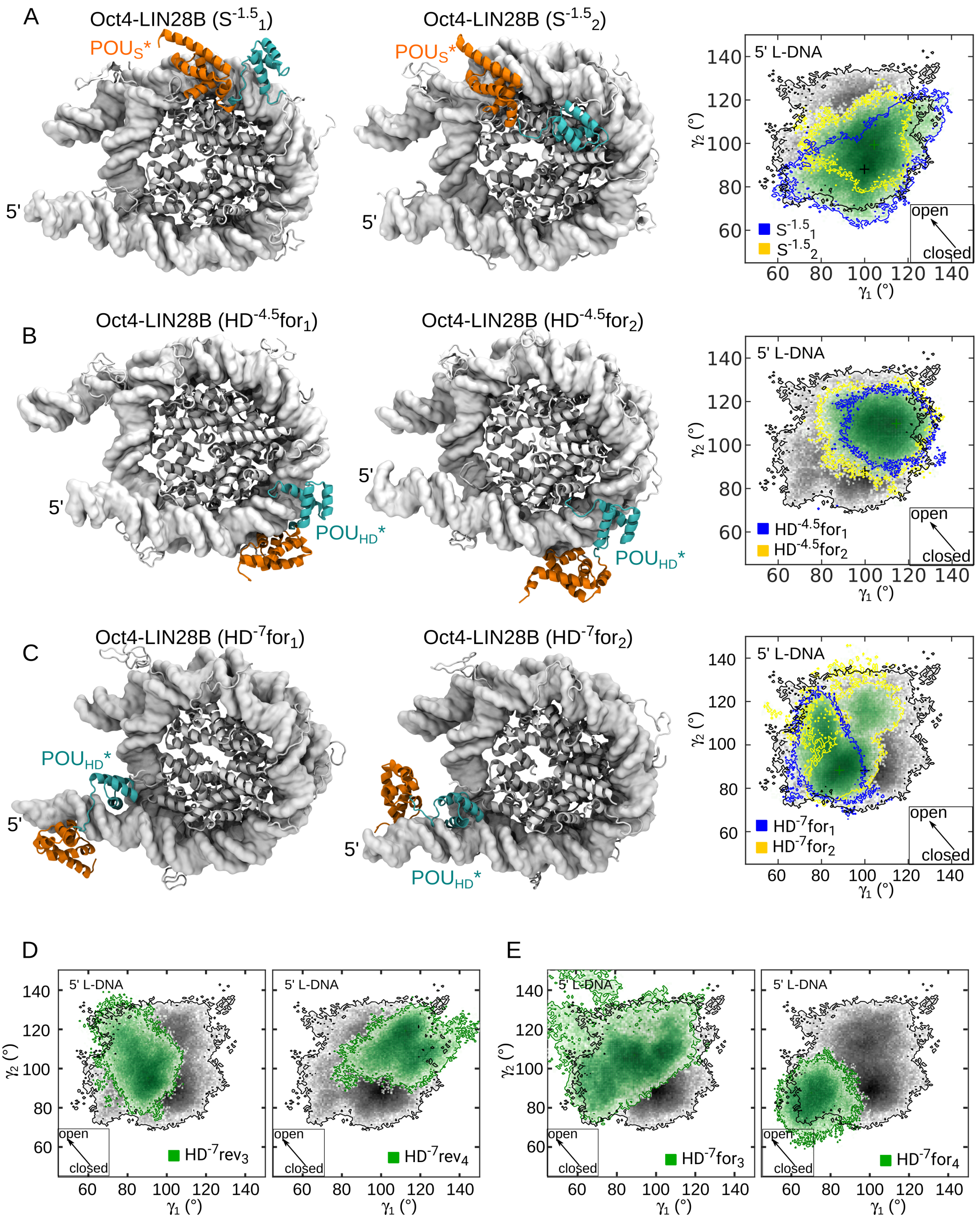


**Figure S4: Breathing motions of LIN28B in simulations of the Oct4-LIN28B complex with Oct4 bound to different sites.** (**A-C**) Representative structures and 2D histograms depicting the sampling of the $\gamma_1/\gamma_2$ conformational space from the simulations of Oct4 bound to LIN28B on the $S^{-1.5}$ site (A), $HD^{-4.5}$ site in forward orientation (B), and $HD^{-7}$ site in forward orientation (C). (**D-E**) 2D $\gamma_1/\gamma_2$ histograms from additional simulations of Oct4 bound to LIN28B on the $HD^{-7}$ site in reversed (D) and forward orientation (E). In the histograms, the sampling from the simulations of the free nucleosome is in black, whereas the sampling from the simulations of Oct4-LIN28B complex is in green. The yellow and blue contours indicate the sampling from the individual 1 $\mu$s simulations. The arrows in the square inserts indicate the direction of the nucleosome opening. The "*" labels the sequence specific bound subdomain. See also Figure 3.

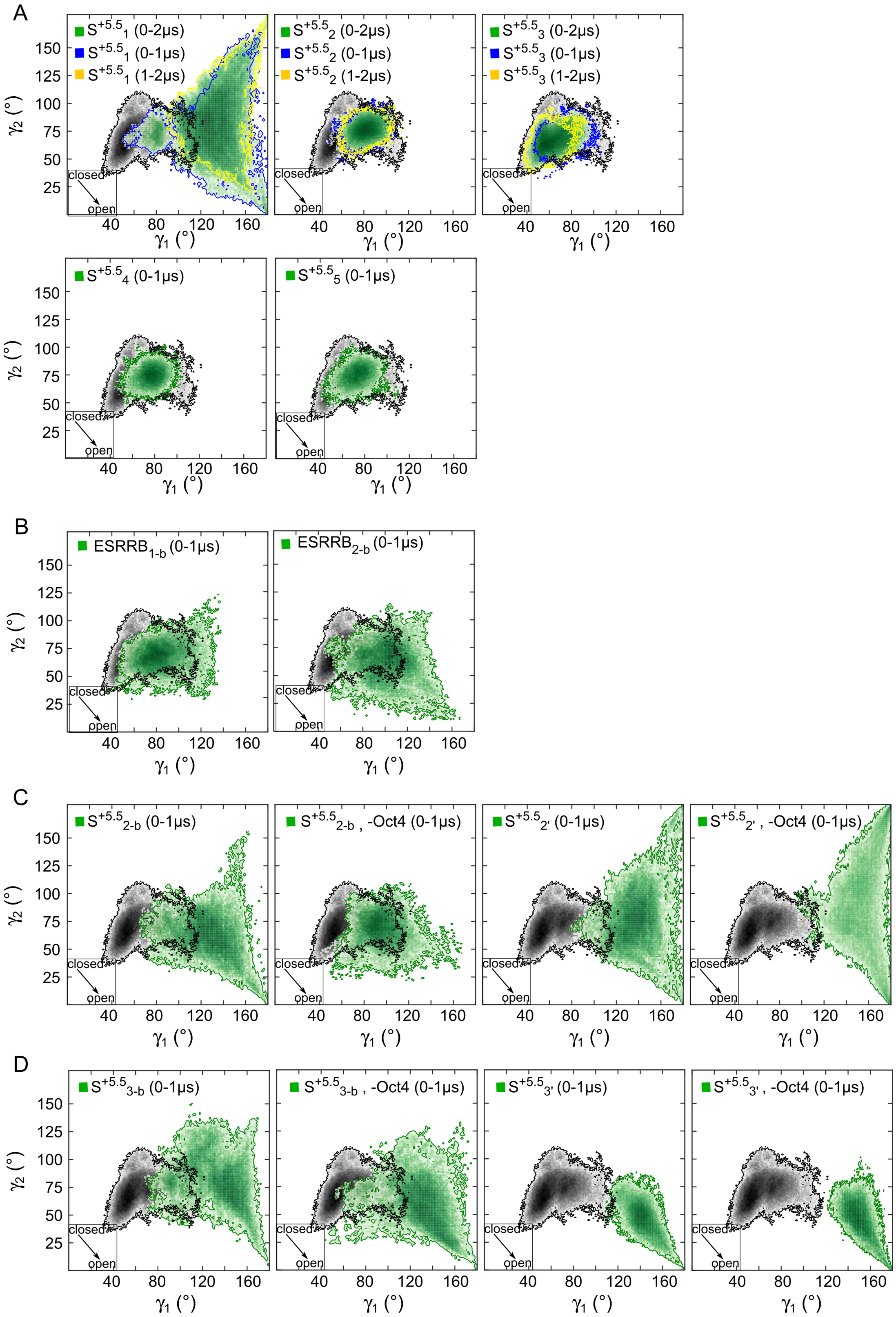


**Figure S5: Breathing motions of ESRRB in classical and biased simulations of the ESRRB alone and the Oct4-ESRRB complex.** (**A-D**) 2D histograms as in Figure S4 depicting the sampling of the $\gamma_1/\gamma_2$ conformational space from: the initial 5 simulations of the Oct4-ESRRB complex (A), the biased simulations of the ESRRB alone (B), the biased simulation ($S^{5.5}{}_{2\text{-b}}$) and the follow-up classical simulation ($S^{5.5}{}_{2'}$) of the Oct4-ESRRB complex started from the $S^{5.5}{}_{2}$ simulation (C), the biased simulation ($S^{5.5}{}_{3\text{-b}}$) and the follow-up classical simulation ($S^{5.5}{}_{3'}$) of the Oct4-ESRRB complex started from the $S^{5.5}{}_{3}$ simulation (D). See also Table 1 and Figure 3.

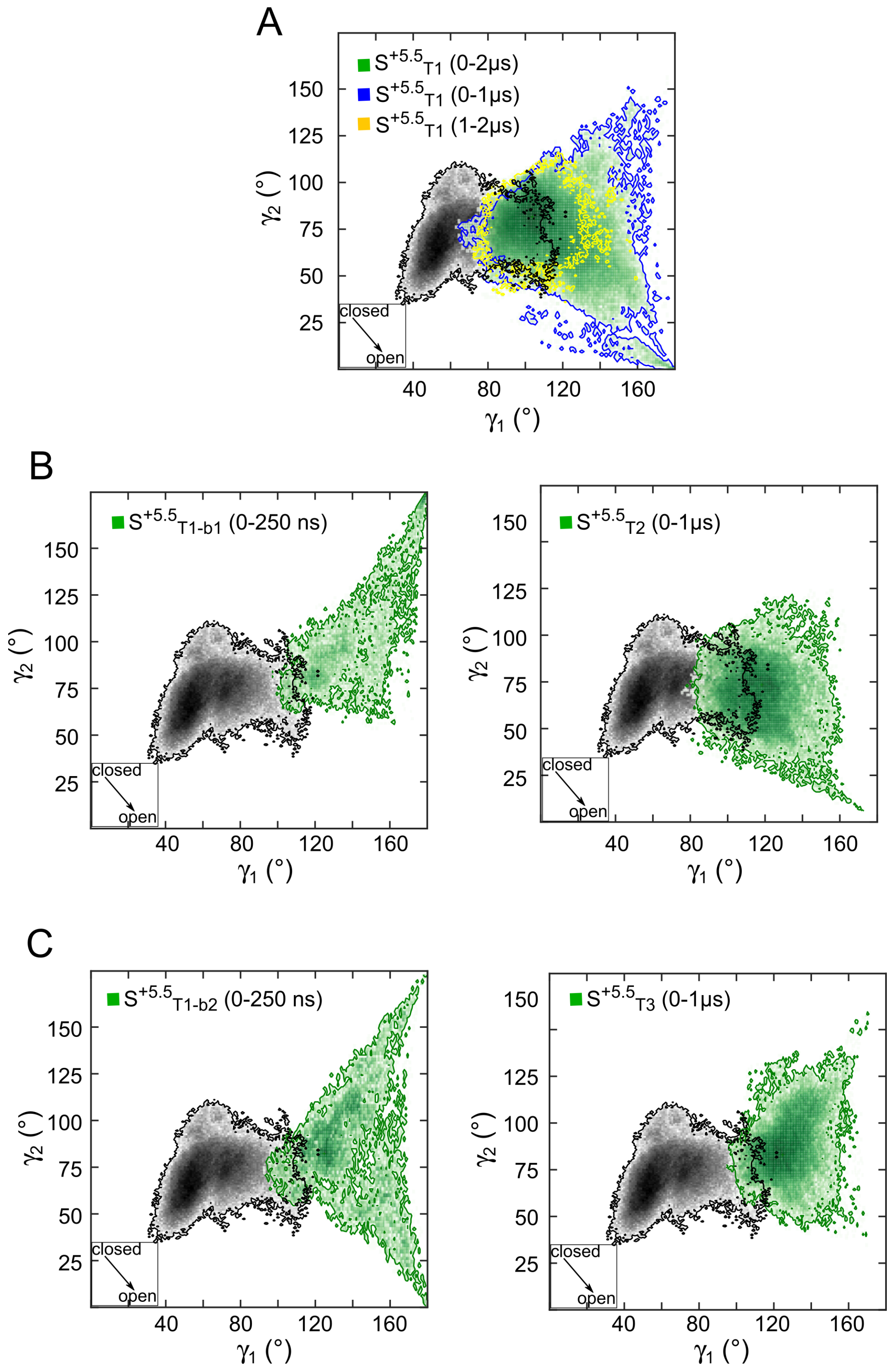


**Figure S6: Breathing motions of ESRRB in classical and biased simulations of the Oct4-ESRRB complex started with modified H3 and H2AC tail configurations.** 2D histograms as in Figures S4, S5 depicting the sampling of the $\gamma_1/\gamma_2$ conformational space from the simulations $\mathrm{S}^{5.5}{}_{\mathrm{T1}}$, $\mathrm{S}^{5.5}{}_{\mathrm{T1\text{-}b1}}$, $\mathrm{S}^{5.5}{}_{\mathrm{T2}}$, $\mathrm{S}^{5.5}{}_{\mathrm{T1\text{-}b2}}$, and $\mathrm{S}^{5.5}{}_{\mathrm{T3}}$ See also Table 1 and Figure 3.

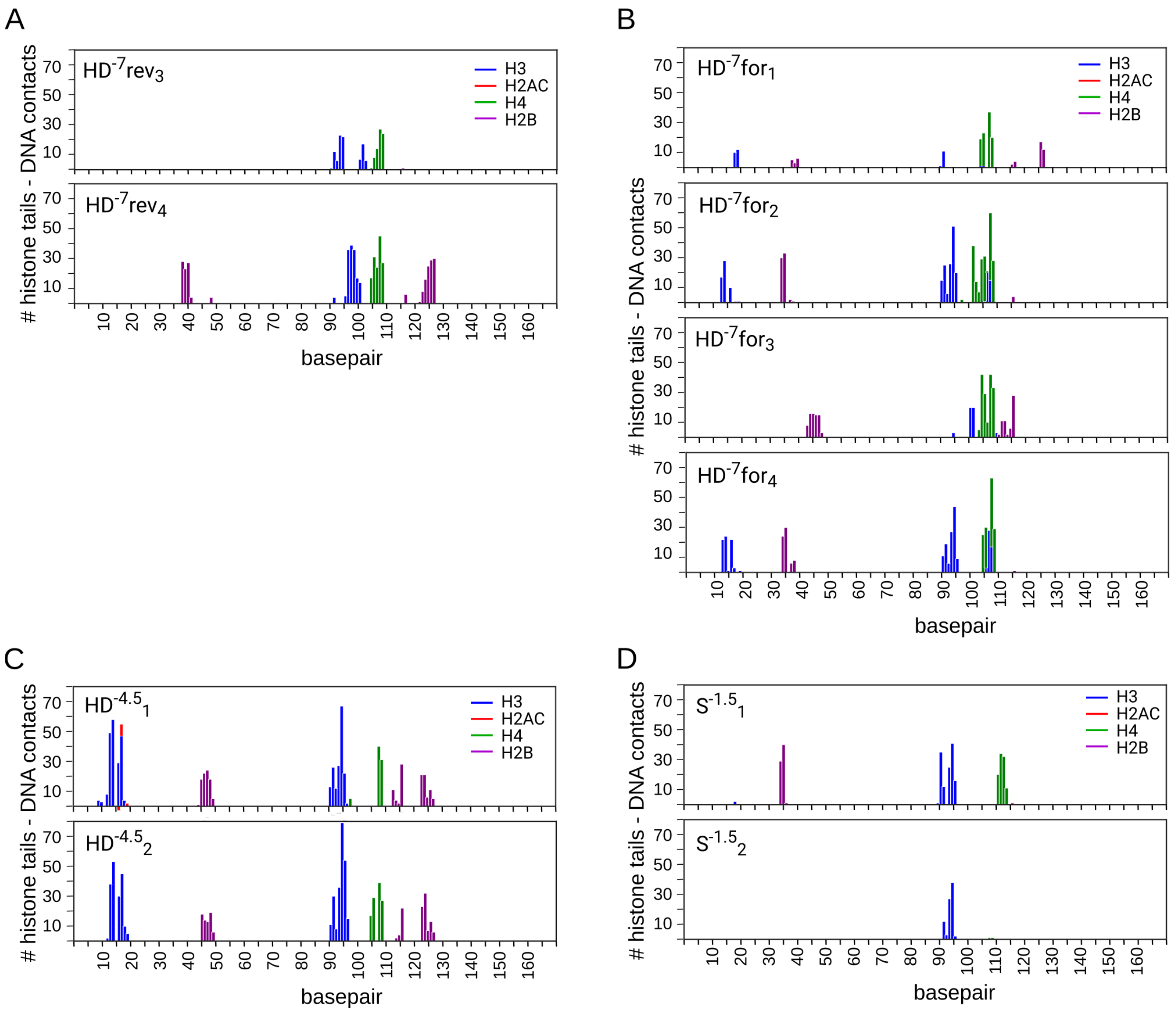


**Figure S7: Histone tail - DNA interaction profiles in the Oct4-LIN28B complex** The number of stable contacts between the histone tails and the DNA is shown, with contacts defined as stable if they are present in more than the 75% of the simulation (as in Figure 4). (**A-B**) Additional simulations with Oct4 bound on the HD$^{-7}$ site, in the reverse (A) and forward (B) orientations. (**C-D**) Simulations with Oct4 bound on the HD$^{-4.5}$ (C) and HD$^{-1.5}$ (D) sites. The data for H3, H2AC, H2B and H4 tails are in blue, red, purple, and green, respectively. See also Tables 1, S1 and Figure 4.

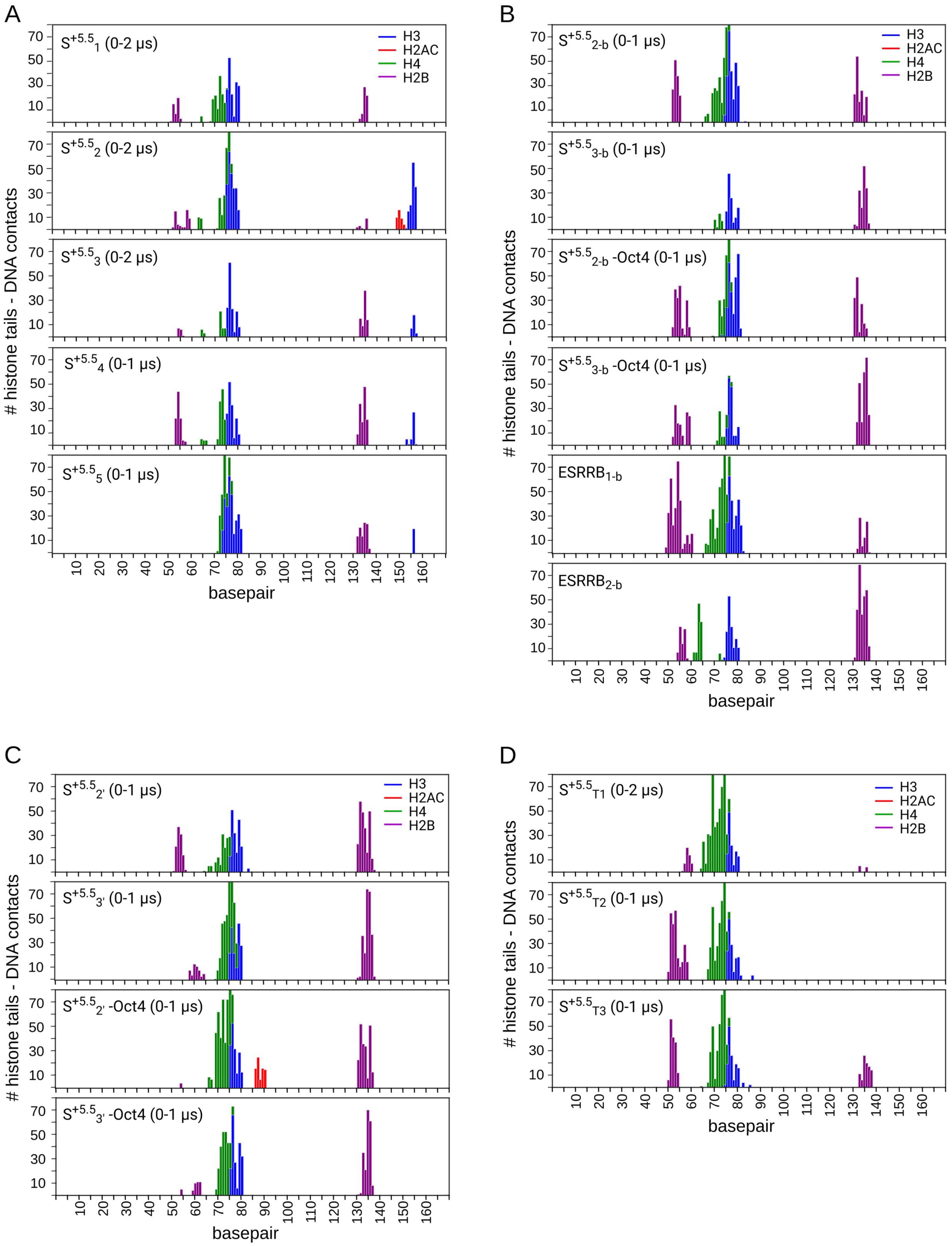


**Figure S8: Histone tail - DNA interaction profiles in the ESRRB and the Oct4-ESRRB complex.** The plots are similar to those in Figure S7. (**A**) The initial 5 classical simulations. (**B**) The biased simulations (**C**) The classical simulations started after the biased simulations from B. (**D**) The classical simulations started with the modified H3 and H3AC tail configurations. See also Table 1, S1 and Figure 4.

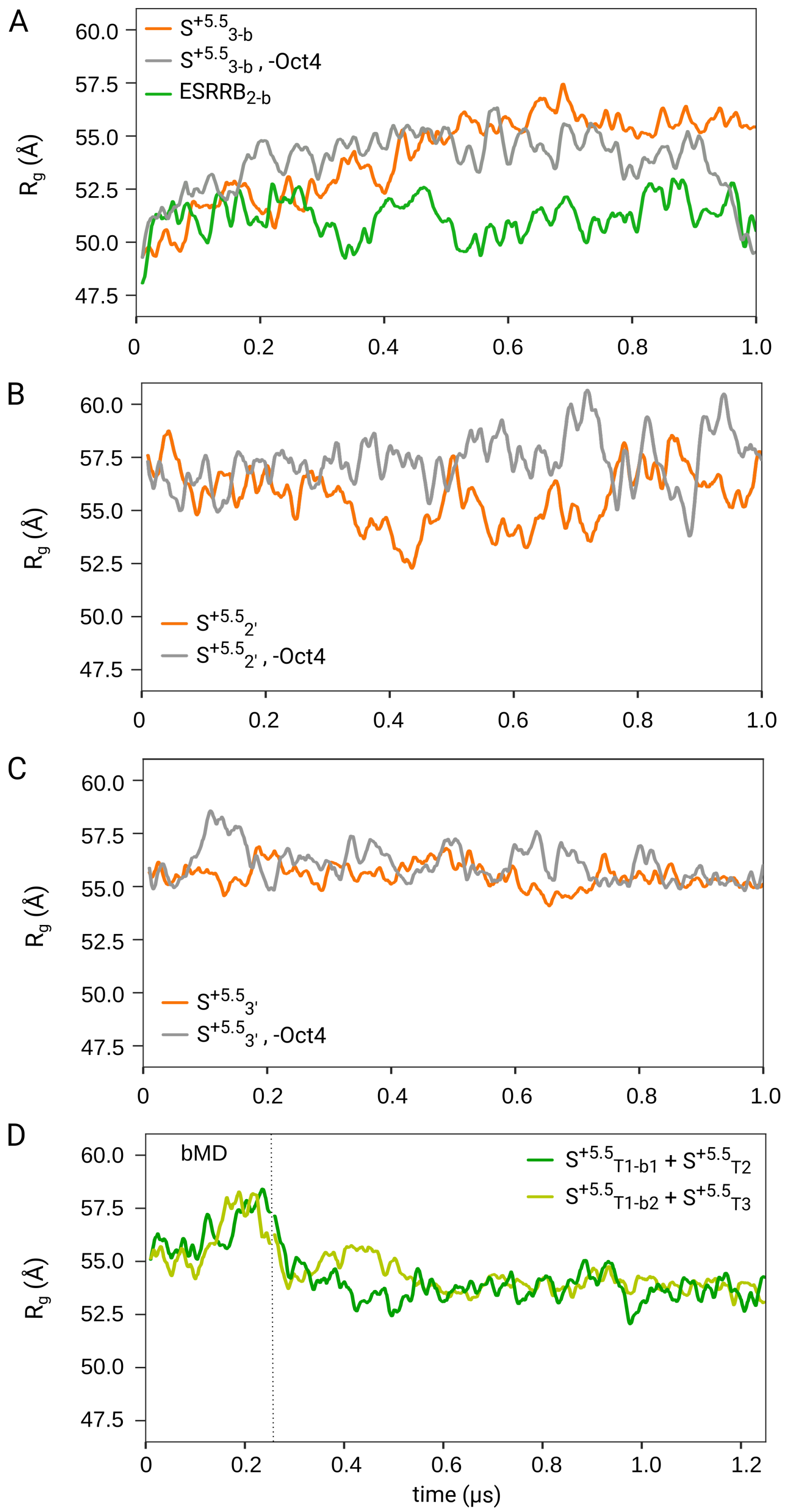


**Figure S9: Nucleosome opening in biased simulations followed by classical simulations of the Oct4-ESRRB complex.** Time series of $R_g$ from additional biased and classical simulations. (**A**) Biased simulations of the ESRRB alone ($ESRRB_{2\text{-b}}$ (green) started from $ESRRB_2$) and of the Oct4-ESRRB started from $S^{5.5}{}_3$ with (orange) and without Oct4 (gray). (**B**) Classical simulations with (orange) and without (gray) Oct4 started from the biased $S^{5.5}{}_{2\text{-b}}$. (**C**) Classical simulations with (orange) and without (gray) Oct4 started from the biased $S^{5.5}{}_{3\text{-b}}$. (**D**) biased and classical simulations started with modified H3 and H2AC tail configurations ($S^{5.5}{}_{T1\text{-b1}}$ followed by $S^{5.5}{}_{T2}$ (dark green) and $S^{5.5}{}_{T1\text{-b2}}$ followed by $S^{5.5}{}_3$ (light green)). See also Table 1, S1 and Figure 5.

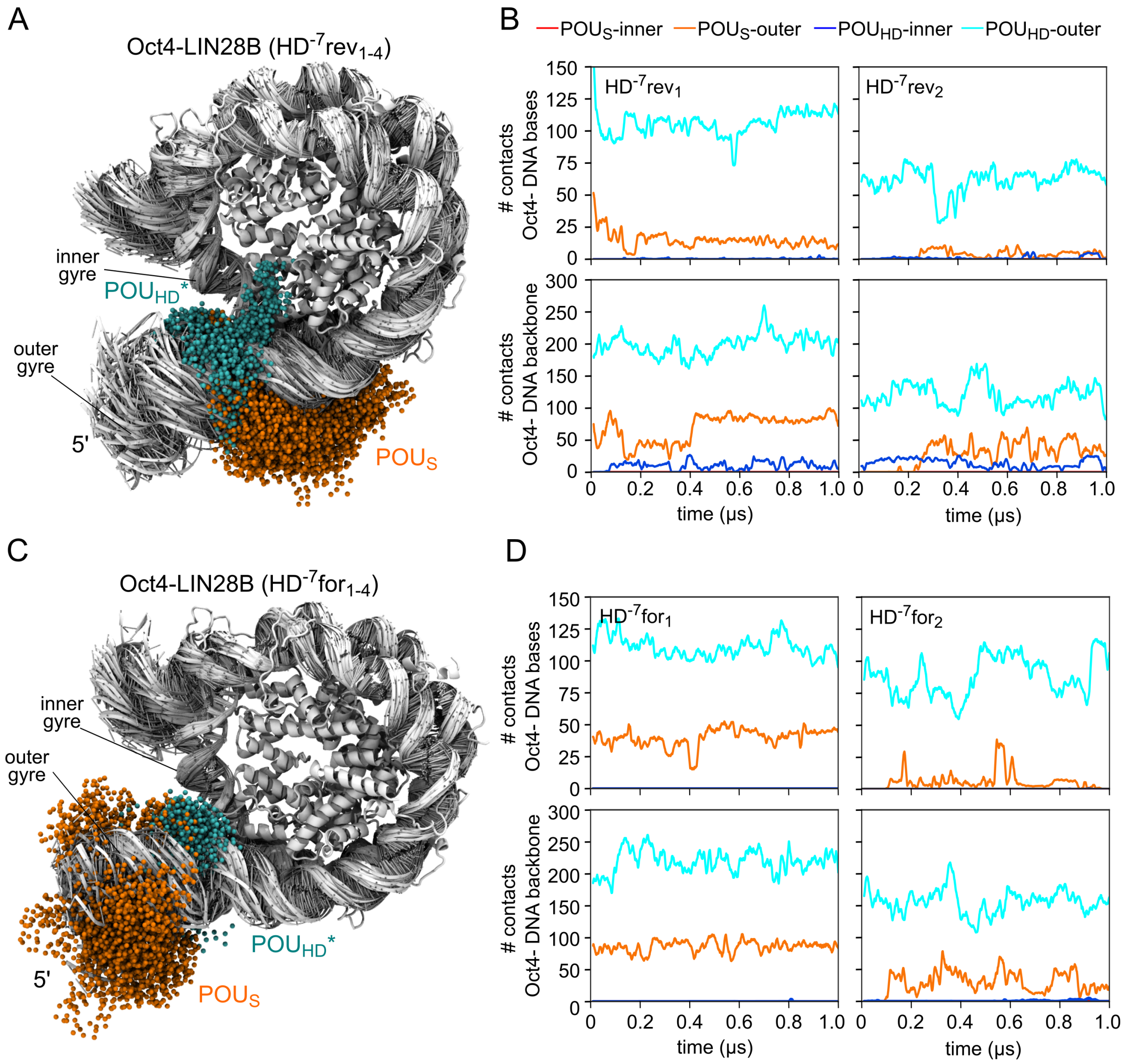


**Figure S10: Sequence specific binding and nonspecific DNA exploration on LIN28B by the two Oct4 subdomains.** (**A**) Nucleosome sampling by the Oct4 subdomains in 4 $\mu$s aggregate simulations time with Oct4 bound to the HD$^{-7}$ site in reverse orientation. (**B**) Number of contacts between the POU$_S$ (red, orange) or the POU$_{HD}$ (blue, cyan) with DNA bases (sequence specific, upper plots) or DNA backbone (non-specific, lower plots) of the inner gyre (red, blue) or the outer gyre (orange, cyan) in two of the simulations shown in (A) (HD$^{-7}$rev$_1$ and HD$^{-7}$rev$_2$). (**A**) Nucleosome sampling by the Oct4 subdomains in 4 $\mu$s aggregate simulations time with Oct4 bound to the HD$^{-7}$ site in forward orientation. (**D**) Same as (B) but in two of the simulations shown in (C) (HD$^{-7}$for$_1$ and HD$^{-7}$for$_2$). See also Tables 1, S1 and Figure 6.
